# Regulation of FXR1 by alternative splicing is required for muscle development and controls liquid-like condensates in muscle cells

**DOI:** 10.1101/818476

**Authors:** Jean A. Smith, Ennessa G. Curry, R. Eric Blue, Christine Roden, Samantha E. R. Dundon, Anthony Rodríguez-Vargas, Danielle C. Jordan, Xiaomin Chen, Shawn M. Lyons, John Crutchley, Paul Anderson, Marko E. Horb, Amy S. Gladfelter, Jimena Giudice

## Abstract

Fragile-X mental retardation autosomal homolog-1 (FXR1) is a muscle-enriched RNA-binding protein. FXR1 depletion is perinatally lethal in mice, *Xenopus,* and zebrafish; however, the mechanisms driving these phenotypes remain unclear. The *FXR1* gene undergoes alternative splicing, producing multiple protein isoforms and mis-splicing has been implicated in disease. Furthermore, mutations that cause frameshifts in muscle-specific isoforms result in congenital multi-minicore myopathy. We observed that *FXR1* alternative splicing is pronounced in the serine and arginine-rich intrinsically-disordered domain; these domains are known to promote biomolecular condensation. Here, we show that tissue-specific splicing of *fxr1* is required for *Xenopus* development and alters the disordered domain of FXR1. FXR1 isoforms vary in the formation of RNA-dependent biomolecular condensates in cells and *in vitro*. This work shows that regulation of tissue-specific splicing can influence FXR1 condensates in muscle development and how mis-splicing promotes disease.

**HIGHLIGHTS:**

- The muscle-specific exon 15 impacts FXR1 functions
- Alternative splicing of *FXR1* is tissue- and developmental stage specific
- FXR1 forms RNA-dependent condensates
- Splicing regulation changes FXR1 condensate properties

## INTRODUCTION

Fragile-X mental retardation autosomal homolog-1 *(FXR1)* and *FXR2* are vertebrate homologues of the Fragile-X mental retardation-1 (*FMR1*) gene. These genes encode three RNA-binding proteins (RBPs) that comprise the Fragile-X (FraX) protein family (Zarnescu and Gregorio, 2013). FraX proteins regulate mRNA transport, stability, and translation (Darnell et al., 2009). While FMR1 is associated with neuronal functions, FXR1 is highly enriched in striated muscles where it localizes to the Z-discs and costameres and is associated with muscle function in multiple organisms (Huot et al., 2005; Mientjes et al., 2004; Van’t Padje et al., 2009; Whitman et al., 2011; Zarnescu and Gregorio, 2013).

Alternative splicing results in several FXR1 isoforms with varying abundances in different cell types (Khandjian et al., 1998; Kirkpatrick et al., 1999). Isoform balance is altered in myotonic dystrophy (Orengo et al., 2008), facioscapulohumeral muscular dystrophy (Davidovic et al., 2008), and diabetes (Nutter et al., 2016) suggesting that *FXR1* pre-mRNA splicing is central to function. Supporting this, mutations causing frameshifts in muscle-specific isoforms are associated with congenital multi-minicore myopathy in humans (Estañ et al., 2019). However, it is unclear whether phenotypes arise from neomorphic frameshifts or loss of muscle-specific protein sequences (Estañ et al., 2019). So far, the phenotypes of FXR1 manipulation have been observed using methods that affect all splice isoforms. Thus, the mechanism relating *FXR1* pre-mRNA splicing to its function in muscle development is not understood. Therefore, we examined the importance of muscle-specific *FXR1* splicing in development.

Muscle-specific FXR1 isoforms contain a longer primary sequence than in other tissues: with a predicted 300 amino acid long intrinsically-disordered domain (IDD) at the C-terminus. Numerous RBPs contain disordered or low-complexity sequences that are associated with biomolecular condensation or liquid-liquid phase separation (LLPS) (Banani et al., 2017). We postulated that alternative splicing may regulate the IDD and thus FXR1 condensation. RNA-rich granules are prominent in large cells such as neurons, where transport granules package mRNAs for local translation (Kiebler and Bassell, 2006), and in multinucleated fungi where they promote local control of the cell cycle and cell polarity (Lee et al., 2013, 2015). We hypothesized that alternative splicing events within the IDD of FXR1 regulate biomolecular condensates for patterning developing muscle.

In this study, we first examined the splicing patterns of *Fxr1* pre-mRNA and observed that blocking the expression of muscle-specific isoforms leads to alterations in *Xenopus* development *in vivo* and in cultured muscle cell differentiation. We further found that FXR1 forms spherical, liquid-like assemblies in both developing myotubes and cultured U2OS cells and more gel-like assemblies *in vitro*. Additionally, both disordered sequences and RNA binding contributed to condensate assembly and different isoforms vary in the properties of the condensates they form. In summary, this study links alternative splicing of FXR1 to LLPS in muscle development and disease.

## RESULTS

### Splicing of *fxr1* exon 15 impacts the development of *Xenopus*

Recessive mutations in muscle-specific isoforms of FXR1 are associated with congenital multi-minicore myopathy in humans (Estañ et al., 2019). These mutations result in a frameshift in *FXR1* transcripts containing exon 15. It is unclear whether multi-minicore myopathy is the result of exon 15 loss, a neomorphic function conferred by the frameshift, or a combination of both. To investigate *FXR1* exon 15 function in development, we removed or mutated this exon in *Xenopus laevis* (*X. laevis*) and *Xenopus tropicalis* (*X. tropicalis*). We chose *Xenopus* because there are only two characterized Fxr1 splice isoforms, which differ solely by the inclusion of exon 15 (Huot et al., 2005). Further, protein sequences of exon 15 from *X. tropicalis* and the two alloalleles of *X. laevis*, 5S and 5L, differ from mouse and human by only one amino acid (**Fig. 1A**).

**Fig. 1.**
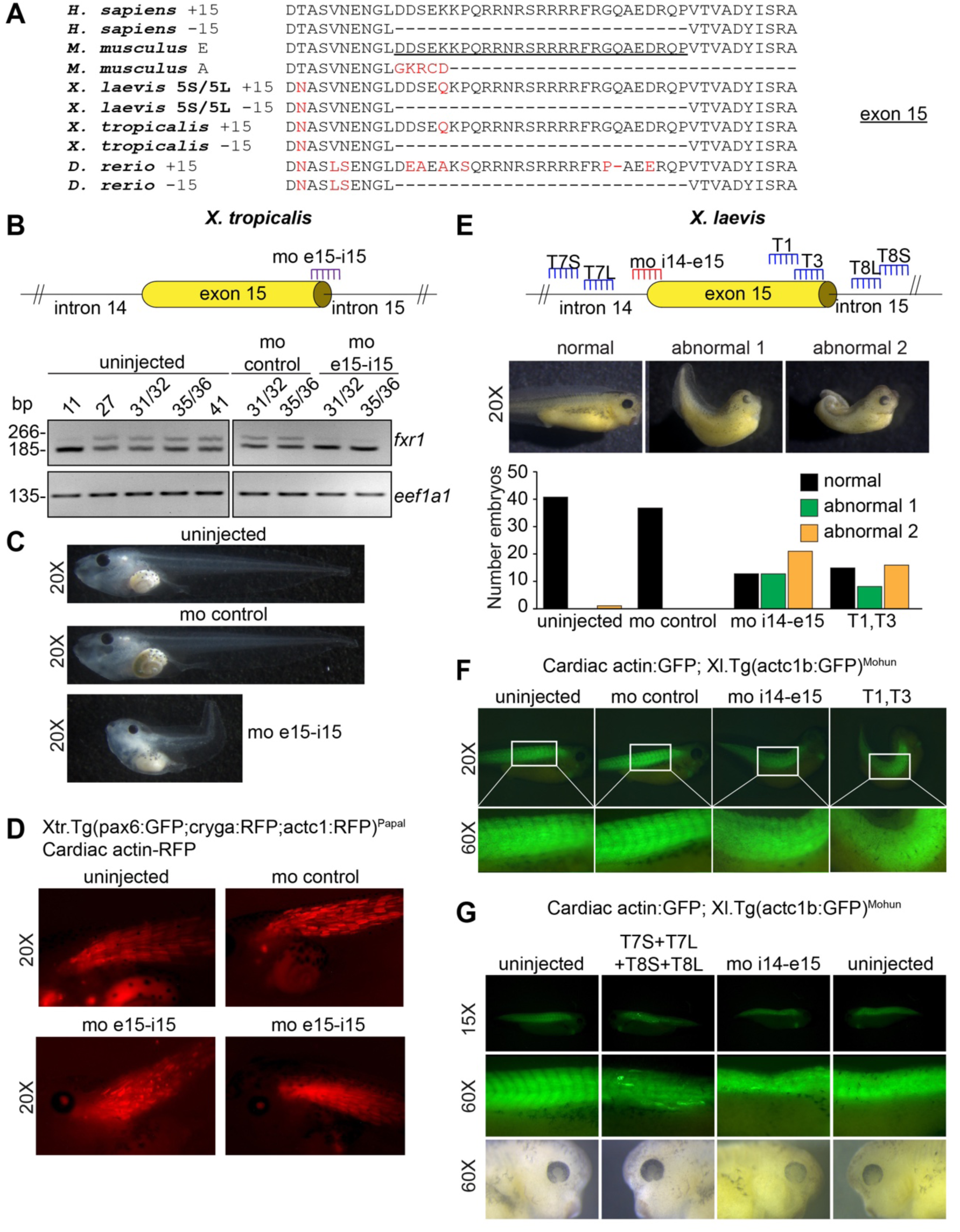
Splicing of *fxr1* exon 15 regulates *Xenopus* development. **A.** Conservation of FXR1 exon 15 amino acid sequences in human, mouse*, X. laevis*, *X. tropicalis*, and zebrafish. Red: deviation relative to mouse/human sequence. Mouse exon 15 is underlined. **B.** A morpholino (MO) targeting exon 15 (mo-e15i15) was used in *X. tropicalis.* Successful blocking of exon 15 inclusion was confirmed by RT-PCR. **C.** Representative images of uninjected embryos or those injected with a control MO (mo-control) or the mo-e15i15 (*X. tropicalis)* at stage 45. **D.** *X. tropicalis* embryos at stage 45. RFP signal marks the somites of uninjected, mo-control, or mo-e15i15 treated embryos. **E.** Position of mo-i14e15 and sgRNAs within the *X. laevis* genome. Representative images and quantification of normal or two abnormal phenotypes (1 and 2) for uninjected, mo-control, mo-i14e15, or sgRNA treated embryos. **F-G.** *X. laevis* embryos at stages 35-38. GFP signal marks the somites of uninjected, injected (mo-control or mo-i14e15) and Cas9 protein injected with sgRNAs targeting exon 15 or intron 14 and 15 for both 5S and 5L. White rectangles indicate the region of the somites magnified in the insets in panel **F**. Figure 1 is linked to **Figure S1**.

To identify the role of *fxr1* exon 15 inclusion in *Xenopus* development, delivery of a morpholino antisense oligonucleotide (MO) was chosen to block the splicing recognition of exon 15 in *X. tropicalis* (**Fig. 1B**) as this species is diploid with a single *fxr1* gene. We injected transgenic *X. tropicalis* embryos where the cardiac actin promoter drives RFP expression to label the developing somites (Hartley et al., 2001) at the one or two cell stage with a MO that binds the junction between *fxr1* exon 15 and intron 15 (mo-e15i15) or control MO (mo-control). Blockage of exon 15 was confirmed by reverse transcription PCR (RT-PCR) (**Fig. 1B**). The mo-e15i15 treated embryos showed a loss of the longer exon 15-containing transcripts compared to embryos injected with the mo-control (**Fig. 1B**). Compared with uninjected or mo-control treated embryos, the mo-e15i15 injected embryos exhibited severe morphological defects including curved tails and defects in somite formation (**Fig. 1C-D**), consistent with Fxr1 role in somite and muscle development in *X. laevis* (Huot et al., 2005).

We next sought to compare the effect of exon 15 loss to mutations that create frameshifts, which would mimic patient mutations. For this, we used two approaches: CRISPR/Cas9 editing of exon 15 and blocking the exon 15 splice site using MOs (**Fig. 1E**). For these experiments, we chose *X. laevis* and used a second MO that targets the intron 14 and exon 15 junction (mo-i14e15) to complement the *X. tropicalis* results. Two sgRNAs were designed to target the 3’ end of exon 15 for both *fxr1.L* and *fxr1.S* genes, denoted T1 and T3 (**Fig. 1E**), creating a frameshift in exon 15-containing transcripts (**Fig. S1**). Compared with uninjected or mo-control treated embryos, mo-i14e15 or sgRNA (T1/T3) injected embryos differed significantly at stage 35-36 of development, displaying similar gross morphological defects (**Fig. 1E**). These alterations included tail curving to varying degrees of severity, closely mimicking those observed in *X. tropicalis* (**Fig. 1B-D**). Remarkably, the observed phenotypes are also similar to the pan-isoform knockdown of *fxr1* in *X. laevis* (Gessert et al., 2010; Huot et al., 2005) suggesting that the majority of Fxr1 function is derived from the isoform containing exon 15.

We next assessed the impact of loss or frameshift of *fxr1* exon 15 on somites using a transgenic line with GFP-labeled somites (Latinki et al., 2002). We imaged GFP signal in stage 35-38 embryos injected with sgRNAs (T1/T3) or mo-i14e15. Compared with control tadpoles, the sgRNA (T1/T3) or mo-i14e15 injected tadpoles exhibited gross abnormalities in somite formation, including failure to segment (**Fig. 1F**), similar to those observed to *fxr1* pan-isoform knockdown (Huot et al., 2005) and our experiments in *X. tropicalis* (**Fig. 1D**). We finally confirmed that deletion of exon 15 using CRISPR/Cas9 editing with four sgRNAs targeting introns 14 (T7L, T7S) and 15 (T8L, T8S) resulted in a similar phenotype (**Fig. 1G**) to that obtained using mo-i14e15 (**Fig. 1F**).

Together our data show that mis-splicing of *fxr1* exon 15 in *Xenopus* development results in somite formation defects similar to those observed when all Fxr1 isoforms were depleted in *X. laevis* (Huot et al., 2005). This suggests that the highly conserved protein sequence of exon 15 is essential for *Xenopus* development, and that most of the function of Fxr1 is derived from exon 15 containing proteins.

### Alternative splicing of *Fxr1* pre-mRNA is tissue- and developmental stage-specific

We next confirmed the splicing patterns observed in *Xenopus* development are conserved in mammals. In mice and humans, the *FXR1* gene contains 17 exons (**Fig. 2A**) and gives rise to multiple protein isoforms (**Fig. 2B**) via alternative splicing (Kirkpatrick et al., 1999). Notably, exon 16 inclusion induces a frameshift that increases protein length and alters amino acid composition (**Fig. 2B**). Inspection of deep RNA-sequencing data from C2C12 cell differentiation (Singh et al., 2014) and mouse heart and skeletal muscle development (Brinegar et al., 2017; Giudice et al., 2014) revealed that exons 15 and 16 are regulated by alternative splicing during myogenesis and between birth and adulthood in striated muscles. Previously, alternative splicing of *Fxr1* pre-mRNA in adult mouse tissues was noted but never quantitatively analyzed (Davidovic et al., 2008; Huot et al., 2005; Kirkpatrick et al., 1999).

**Fig. 2.**
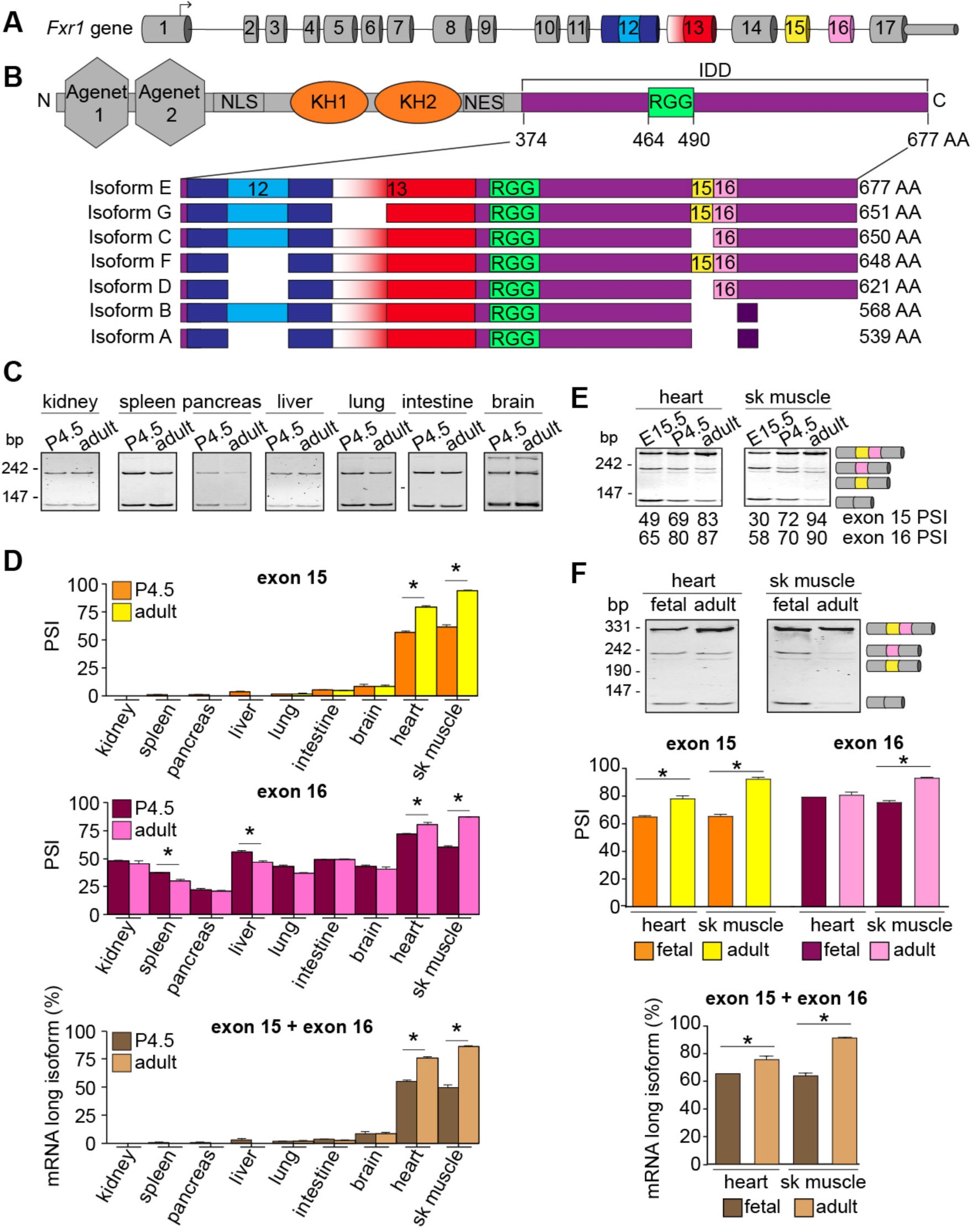
Alternative splicing regulates FXR1 in a tissue- and developmental stage-specific manner. **A.** Mouse *Fxr1* gene structure. **B.** Location of the alternatively spliced region within FXR1. Darker purple in isoforms A and B denotes different amino acids. **C-D.** Splicing of exons 15 and 16 of *Fxr1* pre-mRNA was evaluated by RT-PCR in non striated muscles of neonatal (P4.5) and adult (4 months) mice (**C**). The percent inclusion (PSI) of exon 15 or exon 16 and the abundance of transcripts evaluated by densitometry (**D**). **E.** *Fxr1* pre-mRNA splicing was similarly evaluated in mouse skeletal muscles (sk. muscle) and hearts at embryonic day 15.5 (E15.5), P4.5, and adulthood. **F.** *FXR1* pre-mRNA splicing was similarly evaluated in commercial human RNA samples. Results: mean ± s.e.m. *p≤0.05 (one-way ANOVA test with Bonferroni correction for multiple comparisons), *N*=3-4 (mouse tissues), *N*=2 (fetal human tissues), *N*=3 (adult human tissues). AA: amino acids, bp: base pairs, NES: nuclear export signal, NLS: nuclear localization signal. Figure 2 is linked to **Figure S2**.

Here, we characterized the splicing patterns of exons 12, 13, 15 and 16 and quantitatively measured the percent spliced in (PSI) of each exon (Wang et al., 2008) at different stages of mouse development by RT-PCR. We observed that exon 15 was rarely included in kidney, spleen, pancreas, liver, lung, intestine, or brain at postnatal day 4.5 (P4.5) or adulthood (~4 months) (**Fig. 2C-D**). In contrast, exon 15 was more included in adult hearts and skeletal muscles than in neonates or embryos (**Fig. 2E**). While exon 16 was partially included in non-striated muscle tissues, its inclusion increased during postnatal development only in striated muscles (**Fig. 2C-E**). We further validated that splicing transitions of both exons 15 and 16 are conserved in human heart and skeletal muscle (**Fig. 2F**).

The splicing events in exons 12 and 13 displayed different patterns than exons 15 and 16. All the examined tissues included the entire exon 13 while the insert in exon 12 was mostly skipped in non-muscle tissues (**Fig. S2A-B**). In contrast, in striated muscles almost half the transcripts included the insert (**Fig. S2B**). Unlike exons 15 and 16, the 87-nt insert in exon 12 was not regulated during development in striated muscles.

We then confirmed that the detected splice variants were translated using an antibody against the N-terminus of FXR1 that recognizes all isoforms in Western blots on adult mouse tissue (**Fig. S2C**). Brain and liver expressed a small protein band (~60-65 kDa) corresponding to isoforms A/B. Kidney exhibited the same band as well as an additional low molecular weight band (~45 kDa) (**Fig. S2C**), not corresponding to any known FXR1 isoform (**Fig. 2B**). Heart and skeletal muscle expressed mostly the long FXR1 isoform of ~80 kDa (**Fig. S2C**), corresponding to a combination of isoforms E/F. Overall, these data indicate that *Fxr1* splicing is conserved and leads to the expression of different protein variants in specific tissues and developmental stages.

### C2C12 cell differentiation recapitulates *Fxr1* splicing transitions of muscle development *in vivo*

Given the conserved regulation of *Fxr1* transcripts by splicing we next identified a mammalian cell system, C2C12 cells, where we could examine the functional consequences of splicing. Differentiation of C2C12 myoblasts into myotubes (myogenesis) is a well-established model to study molecular aspects of skeletal muscle biology in culture (Burattini et al., 2004). A previous study implicated FXR1 in muscle cell proliferation (Davidovic et al., 2013) but its role in differentiation was not investigated. By using small interference RNAs (siRNAs), we found that FXR1 depleted cells (**Fig. S3A-B**) formed fewer myotube-like cells than controls (**Fig. S3C**). We further identified that this phenotype was due to a ~50-80% decrease in cell-cell fusion (**Fig. S3D**). Both siRNAs used showed this decrease, with si-*Fxr1*-#2 giving a stronger fusion phenotype and a reduction in the number of nuclei per field of view (**Fig. S3E**). Collectively, these results demonstrate that FXR1 is required for myoblast fusion.

We then systematically quantified the splicing of exons 12, 13, 15 and 16 during myogenesis. Two days after differentiation induction, exons 15 and 16 were highly included (**Fig. 3A**) recapitulating the findings observed in muscle development. Unlike exons 15 and 16, C2C12 cells always include exon 13 (**Fig. 3B**). This is consistent with the above mentioned RNA-sequencing dataset (Singh et al., 2014) and previous reports indicating the alternative 3’ splice site in exon 13 usage is rare (Kirkpatrick et al., 1999). The insert in exon 12 was mostly skipped in myoblasts, while included in ~50% of myotube transcripts (**Fig. 3B**). These findings suggest that myotubes express isoforms E and F at a 1:1 ratio. Finally, we confirmed that during cell differentiation smaller protein isoforms A (539 residues) and B (568 residues) are replaced by the largest protein isoforms E (677 residues) and F (648 residues) (**Fig. 3C**). Collectively, these data suggest that C2C12 cell differentiation reproduces *Fxr1* splicing transitions that take place during striated muscle development *in vivo*, leading us to use this culture model for our molecular studies.

**Fig. 3.**
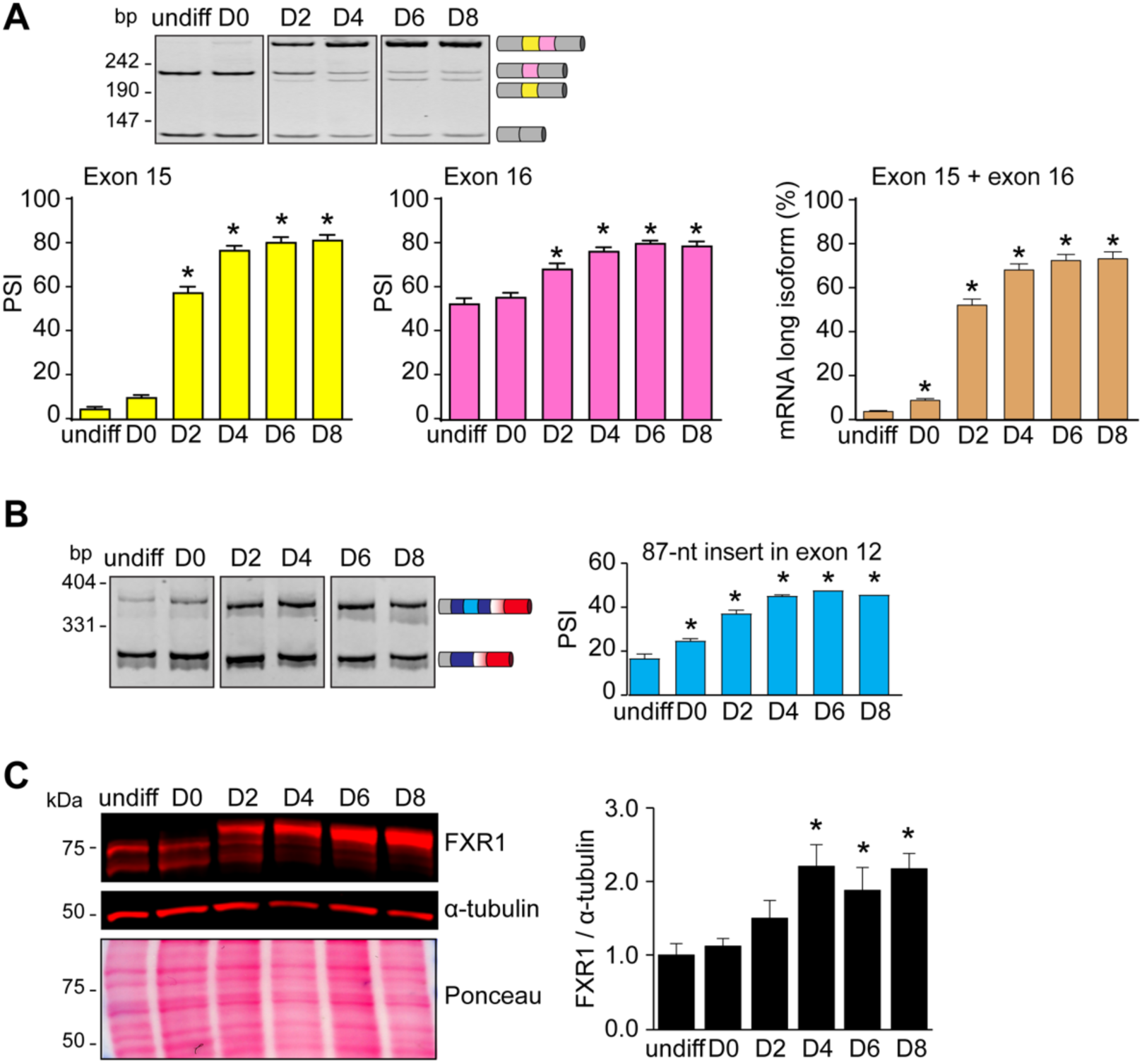
C2C12 cell differentiation reproduces *Fxr1* splicing transitions. RNA or protein was extracted at undifferentiated stage (undiff) and differentiating (D0 to D8) C2C12 cells. **A.** Splicing of exons 15 and 16 was evaluated by RT-PCRs. The PSI was determined by densitometry. **B.** Inclusion or skipping of the 87-nt insert in exon 12 of *Fxr1* pre-mRNA. **C.** Protein lysates were analyzed by Western blotting. Results: mean ± s.e.m. *p≤0.05 *vs* undiff (one-way ANOVA test with Bonferroni correction), *N*=3-5 experiments. bp: base pairs. Figure 3 is linked to **Figure S3**.

### *Fxr1* splicing produces distinct intrinsically-disordered proteins that form condensates

What is the functional consequence of producing splice isoforms of FXR1 with C-terminal extensions in myotubes? To answer this question we examined the disorder tendency the C-termini of isoforms A *versus* E via the IUPRED algorithm (Dosztányi et al., 2005b, 2005a), we found that while the C-termini of both isoforms have IDDs, isoform E has a predicted IDD of ~300 residues, roughly twice as long as isoform A (~160 residues) (**Fig. 4A**). Because a frameshift occurs when exon 16 is included, the different isoform IDDs also vary by amino acid composition; however both are rich in serine and arginine residues (S/R). In SR-rich splicing factors, these regions are subject to post-translational modifications (PTMs), modulating their propensity for disorder (Haynes and Iakoucheva, 2006; Wang et al., 2014; Xiang et al., 2013). Thus, both the IDD length and the potential for PTMs may regulate FXR1 activity during development.

**Fig. 4.**
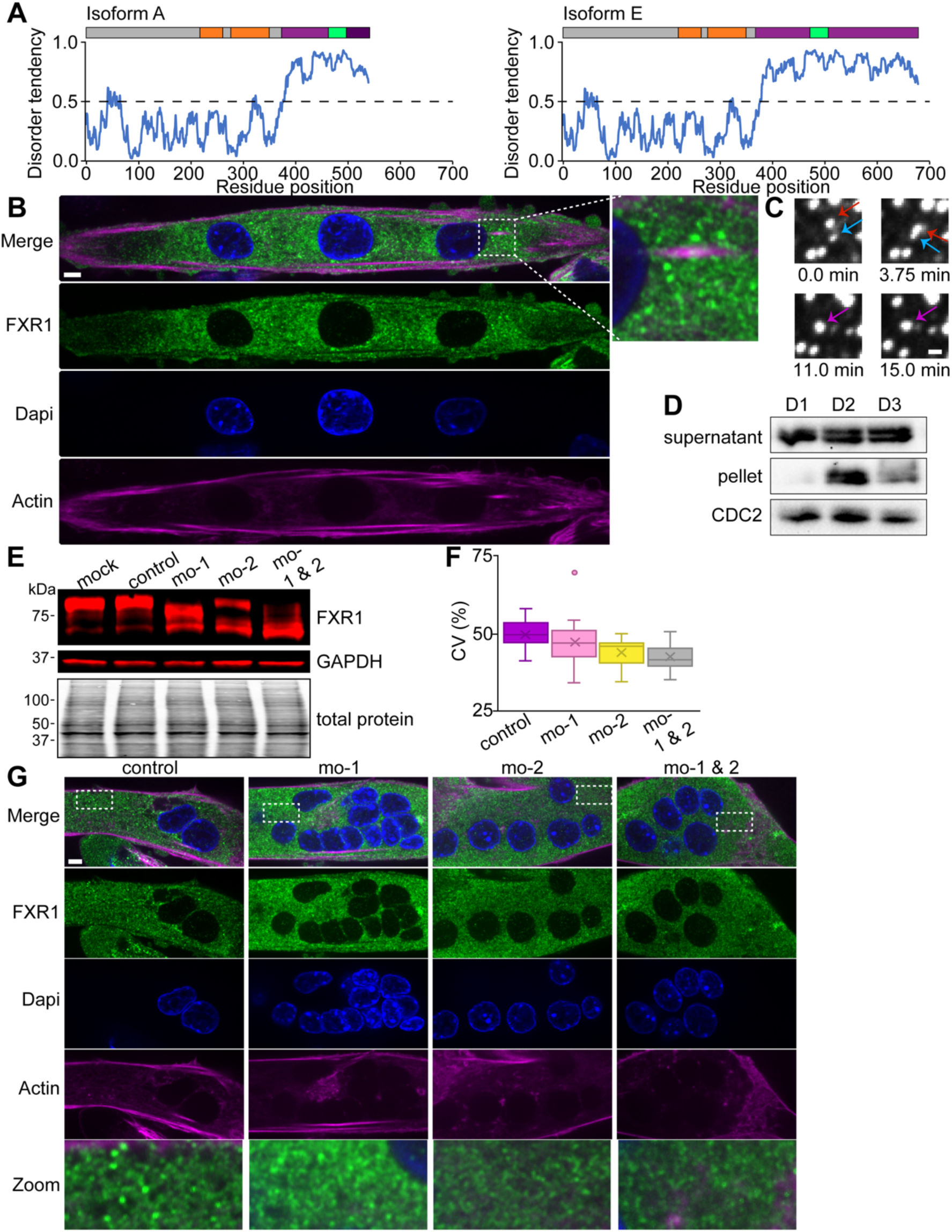
Alternative splicing results in disordered protein isoforms that are capable of phase separation. **A.** Disorder predictions based on the primary amino acid sequence of FXR1 isoforms A and E using the IUPRED algorithm. **B.** Endogenous FXR1 protein was visualized by immunofluorescence in myotubes using an antibody against the N-terminus of FXR1 (recognizes all splice variants). Representative images depicting FXR1 puncta. Scale bar 5 µm. Inset shows droplets magnified 4-fold. **C.** Time-lapse acquisition of human isoform 2 FXR1-eGFP (pAGB828). Two condensates (red and blue arrows) fuse together to form one (purple arrow). Scale bar 1 µm. **D.** C2C12 myoblasts were differentiated for 1, 2, or 3 days (D1, D2, D3). Cell lysates and pellets were analyzed by Western blot using an FXR1 antibody specific to the C-terminus of isoform E. **E-G.** Morpholinos (MOs) targeting exons 15 (mo-1) and/or 16 (mo-2) were delivered in myoblasts and the next day cells were differentiated for 4 days. Cells were analyzed by Western blotting and immunofluorescence using an antibody against FXR1 N-terminus. *Fxr1* splicing was evaluated by RT-PCR (**E**). The coefficient of variation (CV) was calculated as a measurement of FXR1 condensation. p>0.5 mo-1 *vs* control MO, p<0.001 mo-2 and mo1&2 *vs* control MO (**F**). Representative images of MO-treated cells showing FXR1 puncta. Scale bar 5 µm. Bottom row shows droplets magnified 5-fold **(G)**. Results: standard box and whisker plot. Figure 4 is linked to **Figure S4**.

FXR1 is a RBP (Siomi et al., 1995; Zhang et al., 1995), and numerous RBPs contain IDDs that aid in LLPS (Banani et al., 2017; Pak et al., 2016; Uversky et al., 2015; Wang et al., 2014). We therefore hypothesized that FXR1 may form condensates in the myotube cytosol, with isoform E having a higher propensity due to its extended IDD. To determine if FXR1 undergoes LLPS, we visualized FXR1 in C2C12 cells. Immunolabeling of endogenous FXR1 revealed punctate structures throughout the myotube cytosol (**Fig. 4B**). To examine the properties of these structures, *Fxr1* tagged with GFP (*Fxr1-GFP*) was transiently transfected into C2C12 myoblasts that were then differentiated for two days. We observed FXR1-GFP spherical structures in cells that readily fused, suggesting that FXR1 forms liquid-like assemblies (**Fig. 4C**). These droplets were too small and dynamic for partial FRAP studies to accurately assess exchange and rearrangement dynamics. Using an FXR1 antibody against the C-terminus that do not detect isoforms A or B, Western blots were performed on differentiating C2C12 cells (**Fig. 4D**). We found that after two days of differentiation, a portion of FXR1 is visible in the pellet, which may correspond to assembly into insoluble higher-order structures. These data indicate that, like other intrinsically disordered RBPs, FXR1 forms higher-order assemblies and liquid-like droplets in cells.

We next asked if FXR1 condensates are dependent upon alternative splicing. We utilized MOs to redirect endogenous splicing of exon 15 (mo-1) and/or exon 16 (mo-2) in C2C12 cells. MOs were designed to target the splice sites and block exon recognition, thus promoting exon skipping (**Fig. S4A**). MOs were delivered to naïve C2C12 cells that were then differentiated for four days. Both mo-1 and mo-2 blocked exon 15 or 16 inclusion, respectively, (**Fig. S4B**) without altering total mRNA levels (**Fig. S4C**). These MOs also produced smaller protein isoforms excluding these exons (**Fig. 4E**). To determine if altering endogenous *Fxr1* splicing in myotubes caused morphological differences in condensation, we analyzed cells treated with the MOs by immunofluorescence with antibodies against endogenous FXR1. We found that cells lacking exons 15 and 16 formed smaller and fewer FXR1 puncta than control cells expressing the long isoforms. We quantified this effect by measuring the variability in fluorescence intensity of endogenous FXR1 within individual cells and determining the coefficient of variation (CV) of the fluorescent signal (**Fig. 4F-G**). Consistent with having fewer, smaller FXR1-structures, cells with both exons 15 and 16 targeted have the smallest CV, indicative of less heterogeneity in the localization due to more soluble, homogeneously distributed protein (**Fig. 4F-G**). We conclude that alternative splicing of *Fxr1* during myogenesis functions in part to produce isoforms more capable of condensation.

### FXR1 assemblies are protein concentration dependent

If FXR1 is undergoing phase separation to form puncta in muscle cells, then assembly should be protein concentration-dependent. To titrate FXR1 expression, isoform E was placed under the control of a tetracycline (Tet)-inducible promoter in U2OS human osteosarcoma fibroblast cells that had *FXR1*, *FXR2*, and *FMRP* genes knocked out by CRISPR/Cas9 editing (U2OSΔFFF). It was important to use the triple knockout to ensure that endogenous FraX proteins were not acting as seeds for droplets formed with exogenous FXR1. We found that 25 ng/mL Tet-induced expression of mNeonGreen (mNG) tagged-FXR1 isoform E produces approximately endogenous levels (**Fig. 5A**). We hypothesized that isoform E should exhibit increased condensation capacity compared to isoform A because of the data presented in **Fig. 4**. We therefore imaged live U2OSΔFFF cells expressing either mNG-tagged FXR1 isoform A or E at increasing levels of Tet induction (**Fig. 5B**). Cellular condensates were observed at levels below the endogenous protein concentration for both isoforms (**Fig. 5B**). In support of concentration-dependent condensation, the number of cells with droplets increased with Tet concentration (**Fig. 5C**). The proportion of cells with condensates was not statistically different between isoforms A and E. However, we noticed that the condensate morphology differed between the two isoforms (**Fig. 5B**). Isoform A formed large spherical droplets while isoform E formed more irregular shaped bodies. To evaluate droplet morphology, we assigned all the visualized cells to one of the following categories: (1) irregular or non-spherical shaped bodies, (2) small spherical droplets, or (3) large spherical droplets (**Fig. 5D**). Irregular condensates increased in both isoforms with protein concentration with a more pronounced trend for isoform E (**Fig. 5E**). Unexpectedly, isoform A produced more large, spherical droplets. Consistently, both isoforms were enriched in the pellet with increasing Tet concentrations (**Fig. 5F-G**). Despite having a shorter IDD, a higher proportion of isoform A was detected in the pellet at all Tet concentrations (**Fig. 5F-G**) suggesting the composition rather than length of the IDD is important for the traits of the condensates. We suspect that there may be specific PTM differences between myotubes and U2OS cells that could explain the differences between A and E between in these two contexts. Nevertheless, droplet formation is concentration dependent consistent with a phase-separated assembly, morphology and droplets properties.

**Fig. 5.**
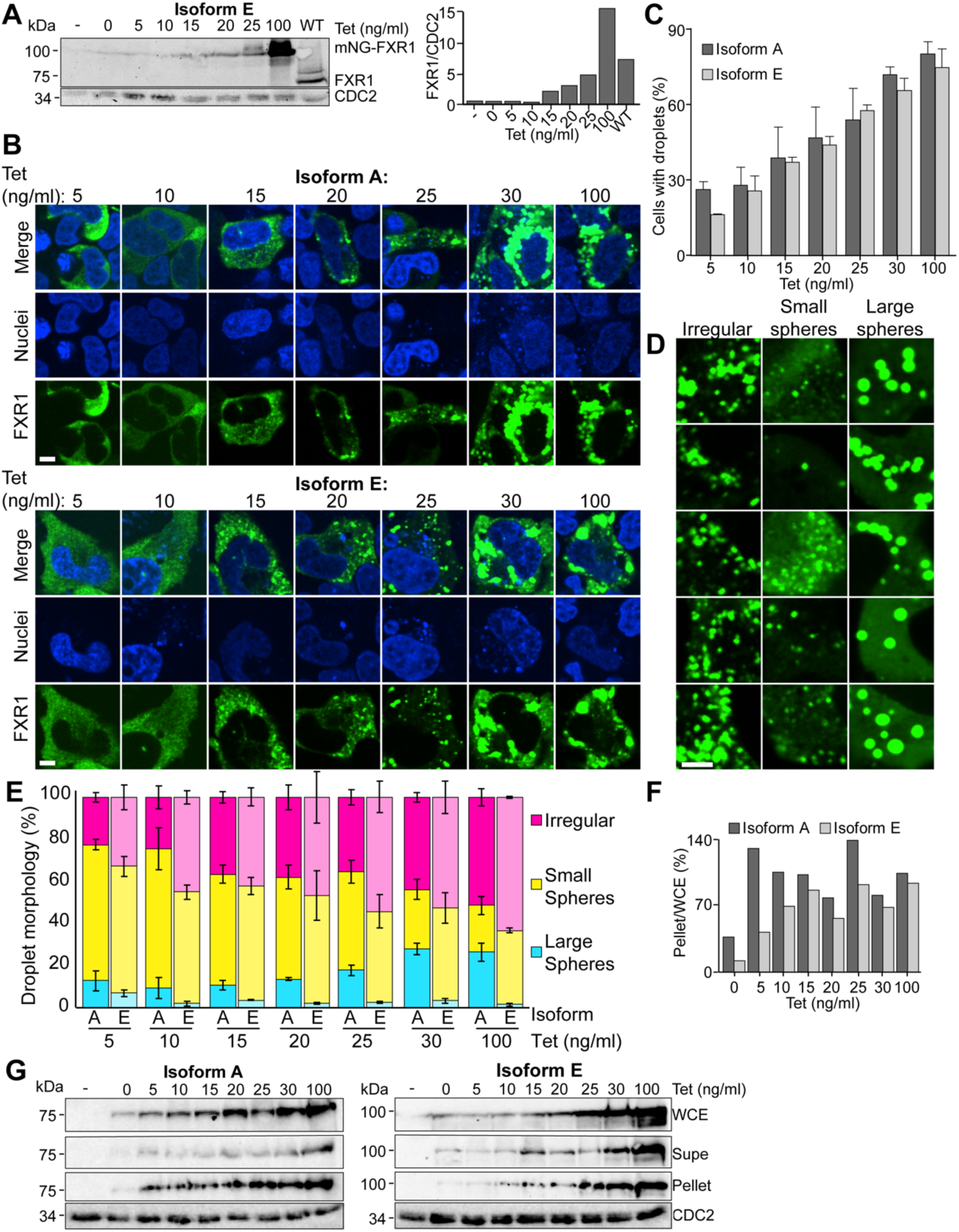
FXR1 phase separation is concentration-dependent. **A.** Whole cell extracts (WCE) from wild type U2OS cells or U2OSΔFFF cells transiently transfected with either an empty vector (pAGB1139) or *P_CMV_2xTetO_2_-mNeonGreen-FXR1 isoform E* (pAGB1162) were prepared. Extracts were analyzed by Western blot. Right panel shows quantification of protein levels normalized to a loading control. **B.** U2OΔFFF cells transiently transfected with an empty vector (pAGB1139), *P_CMV_2xTetO_2_-mNeonGreen-FXR1 isoform A* (pAGB1189), or isoform E (pAGB1162) were lysed. The cell lysate and pellet were analyzed by Western blot as above. The percent of protein in the pellet was quantified as the amount in the pellet out of the amount in the WCE. **C.** Representative images of cells used in panel B at increasing tetracycline (Tet) concentrations. Scale bar 5 µm. **D.** Quantification of droplet formation for individual cells. *n*>720 cells from three independent experiments. **E.** Zoomed in images showing examples of irregular droplets, small spheres, and large spheres. Scale bar 5 µm. **F.** Quantification of droplet morphology. *n*>185 cells from three experiments. Results: mean ± s.e.m.

We next examined the propensity of FXR1 to phase separate *in vitro*. To confirm RNA dependence and isoform specific differences of FXR1 for droplet formation *in vitro*, we purified mouse FXR1 isoforms E and A recombinately from BL21 cells using a 6X HIS GB1 tag (Zhang et al., 2015). GB1 was essential to keep both isoforms A and E soluble during the purification. We observed that in the presence of luciferase RNA, both isoforms E and A had the tendency to slowly aggregate (**Fig. 6A**, **Fig. S5A**) rather than form the droplets as observed *in vivo* (**Figs. 4–5**). Luciferase RNA was chosen as it has previously been shown to induce phase separation of FMR1, which has a similar RNA binding domain as FXR1 (Tsang et al., 2019). There were subtle differences in the morphology of the aggregates between isoforms E and A. Isoform E formed larger aggregates at lower concentration (2 µM) than isoform A which is consistent with isoform E containing a longer disordered sequence. Thus, RNA promotes higher-order assemblies of FXR1 *in vitro*.

**Fig. 6.**
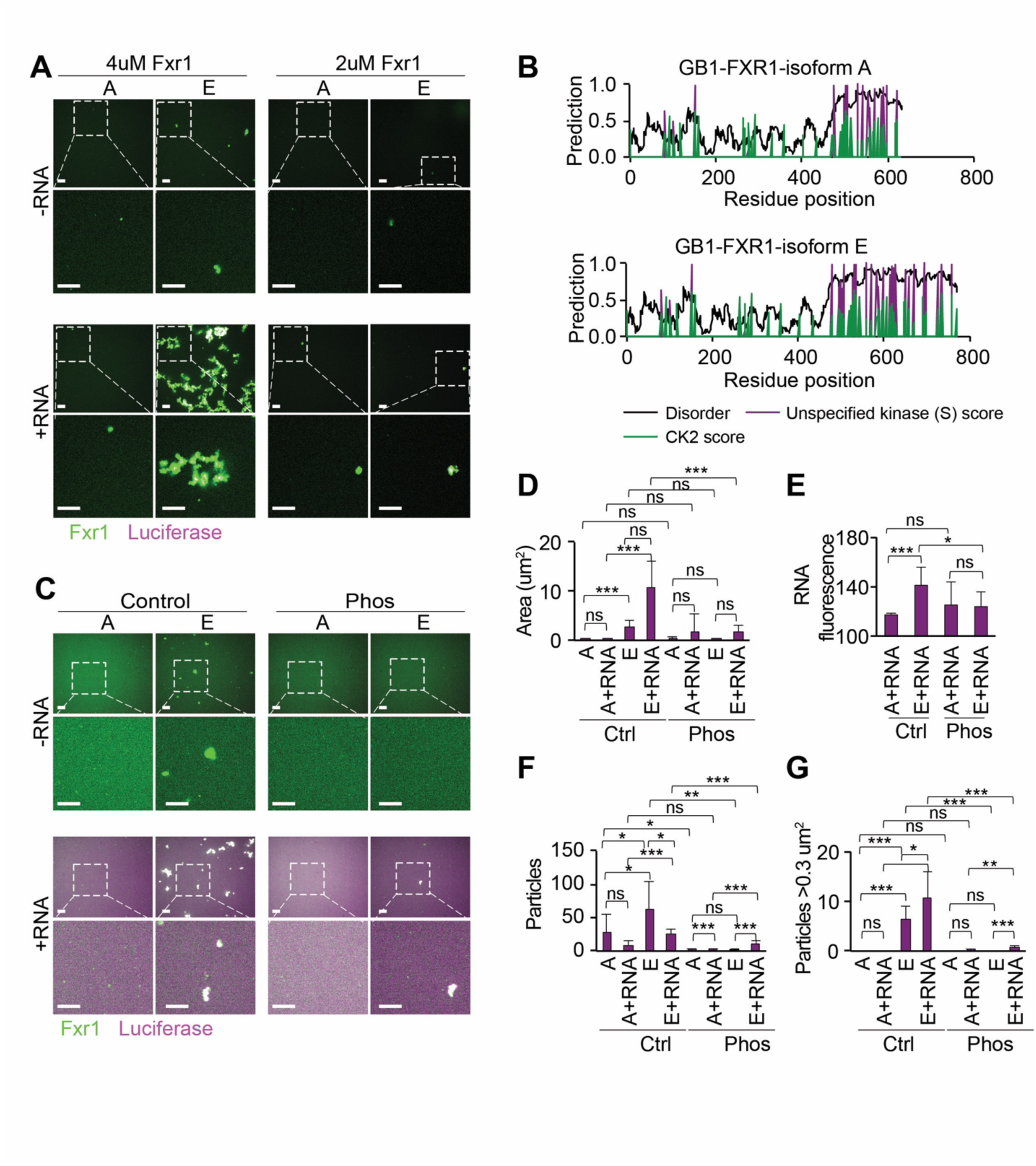
RNA promotes and phosphorylation inhibits FXR1 association *in vitro*. **A.** Representative images of FXR1 isoforms A and E aggregates (2 µM or 4 µM) in the presence or absence of luciferase RNA (magenta). Scale bars 10 µm. **B.** Predicted disorder (IUPRED), unspecified kinase score and CK2 kinase scores for serine are marked for each amino acid. **C.** Representative images of isoform A and E aggregates with or without RNA and with or without phosphorylation. Scale bars 10 µm. **D-G.** Quantification of particle area (**D**), RNA fluorescence within particles (E), number of particles (**F**), and number of particles >0.3 µm^2^ using the images represented in panel **C**. ***p<0.001, **p<0.01, *p<0.05, ns: not significant.

We postulated that a PTM not present in the bacteria was responsible for inducing the droplet (rather than aggregate) behavior in mammalian cells. To this end, we characterized potential phosphorylation sites in FXR1 sequences. Strikingly, a number of serine phosphorylation sites were predicted in the IDD of isoforms A and E, with more sites in the longer IDD of isoform E (**Fig. 6B-C**). Interestingly, a similar concentration of phosphorylation sites in the IDD was not observed in the FMR1 sequence (**Fig. S5B**). To test the importance of FXR1 phosphorylation *in vitro*, we treated isoforms A and E with casein kinase 2 (CK2) either in the presence or absence of ATP and then incubated the resulting protein either in the presence or absence of luciferase RNA. We confirmed that phosphorylation induced a gel shift in FXR1 protein bands (**Fig. S5C**) with a more significant shift in size for isoform E, which had more predicted sites for CK2 phosphorylation (**Fig. 6B**).

Consistent with our previous results, RNA promoted aggregation in both control samples of isoform A and E (**Fig. S5D**). However, phosphorylation tended to reduce (isoform E) or block entirely (isoform A) the aggregation of FXR1 (**Fig. 6D**, **Fig. S5D**). Following 24 h of incubation with RNA, control treated isoform E formed larger aggregates than isoform A (**Fig. 6E**) whereas, phosphorylation blocked aggregate formation for isoform A or substantially reduced aggregation for isoform E (**Fig. 6D**). Phosphorylation had a minimal but significant effect in the observed RNA signal within the particles at 24 h likely due to the reduced large particle formation (**Fig. 6G-H**). This observation is consistent with few predicted phosphorylation sites within the RNA binding domains of FXR1 (N-terminus) shared between isoforms E and A (**Fig. 6F**). Taken together, our studies revealed that at the same protein concentration *in vitro*, isoform E is more prone to form aggregates than isoform A, that RNA accelerates and increases the size of aggregates formed by both isoforms, and that phosphorylation reduces aggregation of both isoforms A and E. This evidence suggests that PTMs such as phosphorylation are critical to the regulation of FXR1 functions *in vivo*.

### FXR1 assemblies are RNA-binding domain dependent

The *in vitro* data suggest RNA is critical for the formation of FXR1 assemblies. To determine which FXR1 domains are required for condensation, we created mutations in the KH domains (*Fxr1^KH1^*, *Fxr1^KH2^*, *Fxr1^KH1/2^*), the RGG box (*Fxr1*^Δ^*^RGG^*), and both a partial truncation (*Fxr1^1-490^*) and complete truncation of the IDD (*Fxr1^1-374^*) of the *Fxr1* coding sequence in the Tet-inducible mNeonGreen-tagged isoform E construct (**Fig. 7A**). We then imaged live U2OSΔFFF cells transiently transfected with each mutant induced to endogenous levels, and quantified the proportion of cells with droplets (**Fig. 7B-C**). When the three RNA-binding domains were mutated (*Fxr1*^Δ^*^RGG KH1/2^*), no condensates were detected. Intermediate phenotypes were observed for the single RNA-binding mutations, implying that the RNA-binding domains are partially redundant for LLPS. All of the five RNA-binding deficient constructs had droplet formation significantly different from wild type.

**Fig. 7.**
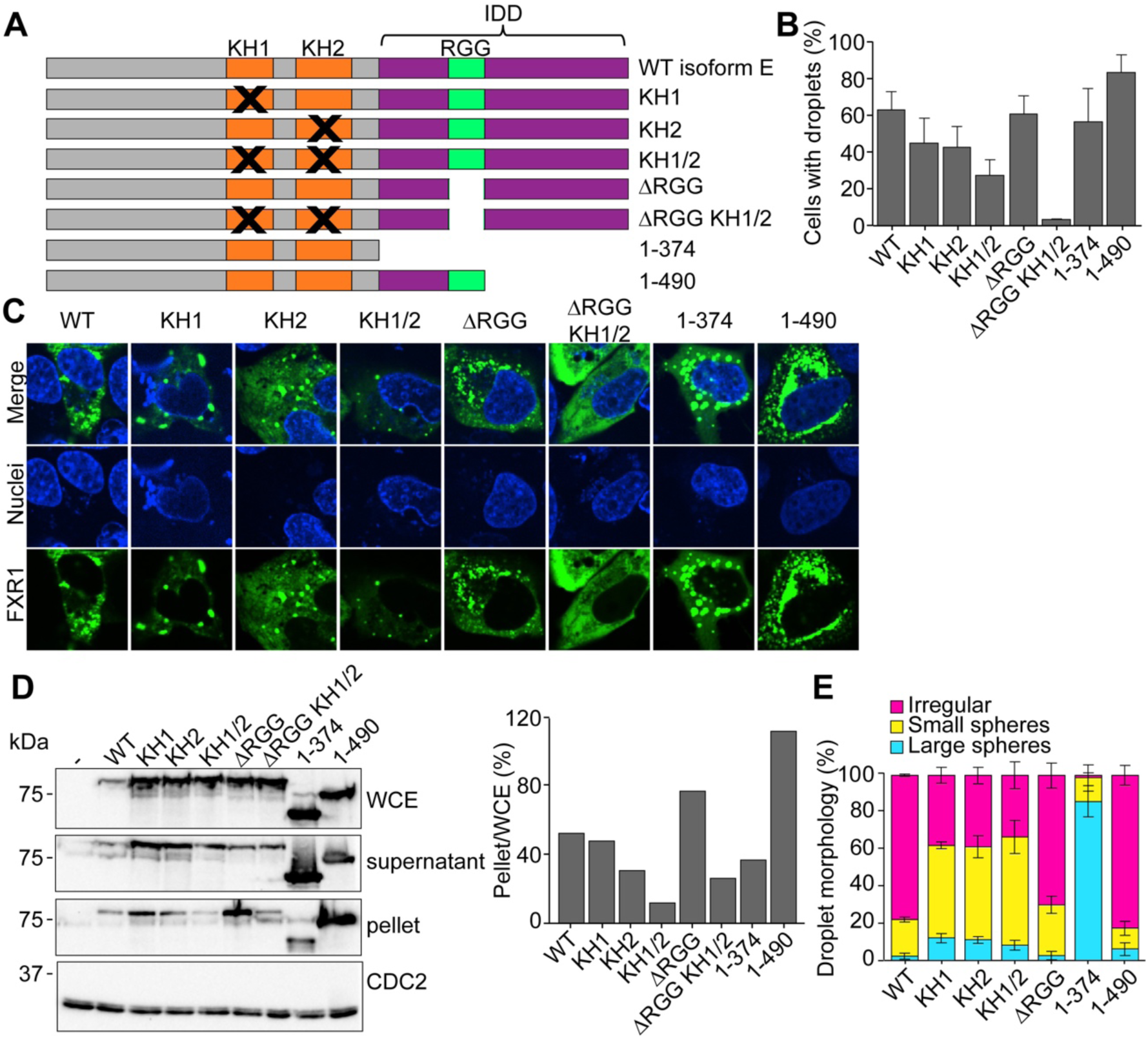
Phase separation is RNA-binding dependent. **A.** Location of mutations in the domains of isoform E. **B-C** U2OSΔFFF cells were transfected with wild type *P_CMV_2xTetO_2_-mNeonGreen-FXR1 isoform E* (pAGB1162), plasmids with mutated KH1 (pAGB1171), KH2 (pAGB1172), KH1/2 (pAGB1173), RGG (pAGB1174), RGG and KH1/2 (pAGB1175), complete IDD (pAGB1176), or partial IDD (pAGB1177). All constructs were induced with 25 ng/mL tetracycline (Tet) to endogenous levels. **B.** Quantification of droplet formation. *n*>350 cells from three experiments. **C.** Representative images of cells expressing the different mutants. Scale bar 5 µm **D.** Whole cell extracts (WCE), lysate, and pellets were analyzed by Western blotting using an antibody against FXR1. The right panel shows the percent protein in the pellet quantified as in Fig. 5. **E.** Quantification of droplet morphology utilizing the same three categories that were used in Fig. 5. Results: mean ± s.e.m.

We next investigated the role of the IDD in droplet formation. When the IDD was truncated to the RGG (*Fxr1^1-490^*) we observed significantly more cells with droplets than wild type, implying that the longer IDD may limit LLPS nucleation. Truncation of the entire IDD (*Fxr1^1-374^*) did not abolish droplet formation, suggesting that the disordered sequence is not essential for phase separation even in cells with no wild type FraX proteins (**Fig. 7B-C**). These data show that the relationship between IDD length and FXR1 droplet formation is non-linear. The whole-cell extract from transfected cells showed that none of the mutant proteins have reduced expression (**Fig. 7D**). Western blots comparing the lysate to the pellet confirmed our observations of condensation from microscopy, with FXR1^ΔRGG^ and FXR1^1-490^ showing the most protein in the pellet, and FXR1^KH1/2^ and FXR1^Δ^ showing the least. While the morphologies of FXR1^KH1/2^ and FXR1^Δ^ are different (**Fig. 7C**), they show similar results in the Western blot assay (**Fig. 7D**).

Droplet morphologies were also affected by some of these mutations. Because the *Fxr1*^Δ^ mutation abolished droplet formation, we did not include this mutant in these analyses. Isoform E showed mostly irregular droplets with some small spherical droplets (**Fig. 7E**), suggesting that small droplets are coalescing incompletely into irregularly-shaped assemblies. Mutation of the KH domains individually or together increased the percentage of cells with small spherical droplets. Without the KH domains, RNA-binding is presumably mediated through the RGG box, which may associate with a different set of RNAs. These RGG-mediated associations still enable small droplets to form but they have limited or slowed fusion. Deletion of the RGG box alone did not alter droplet formation or morphology, implying that the KH domains are the major RNA-binding domains for phase separation. Interestingly, truncation of the IDD back to the RGG box (*Fxr1^1-490^*) did not significantly alter droplet morphology, while deletion of the entire IDD (*Fxr1^1-374^*) caused most cells to have large spherical droplets. This is surprising given that IDDs in other proteins are crucial for this phenotype. These results raise the possibility that RNA interactions are more critical in driving the phase separation of FXR1 than the IDD and that the IDD primarily impacts the material properties of the condensates.

## DISCUSSION

FXR1 is hypothesized to regulate mRNA translation, localization, and stability (Cook et al., 2014; Davidovic et al., 2013; Majumder et al., 2016; Mientjes et al., 2004; Patzlaff et al., 2017; Vasudevan and Steitz, 2007; Zarnescu and Gregorio, 2013); however, there is little known about the mechanisms of function of FXR1 in physiological or pathological contexts. Here, we show quantitative differences in FXR1 isoforms arising from tissue-specific and developmentally regulated alternative splicing and a functional role for *fxr1* splicing in myotube formation and *Xenopus* development. Alternative splicing of *FXR1* pre-mRNA leads to the expression of distinct protein isoforms that vary substantially in the length of the C-terminal IDD. FXR1 can condense into liquid-like droplets in an RNA dependent manner. We observed that the different splice isoforms vary in droplet morphology. These data provide a role for developmentally controlled splicing in regulating the formation of biomolecular condensates.

We hypothesized that FXR1 isoforms expressed in myotubes allow for local translation of large cytoskeleton proteins which are important for proper function of muscle cells. We did observe FXR1 condensation in myotubes, and this condensation was reduced when we blocked the alternative splicing required to form the extended IDD (**Fig. 4**). However, condensates were still observed for the shorter isoform A and for morpholino treated cells which shortens IDDs. This suggests that the change in protein sequence via splicing is important for regulation of condensation with RNA not for confering the ability to condense. We postulate that differential IDD sequences can regulate condensation through PTMs, altered RNA targets, or protein-protein interactions.

FXR1 requires RNA to form droplets such that elimination of all RNA-binding domains blocks LLPS (**Fig. 6**). The RNA-binding sites are partially redundant. RNA drives the phase separation of other RBPs that undergo LLPS including WHI3, LAF-1, and multiple P-granule proteins (Elbaum-Garfinkle et al., 2015; Langdon et al., 2018; Smith et al., 2016; Zhang et al., 2015). Our *in vitro* experiments confirm that luciferase RNA promotes FXR1 aggregation and future experimentations will be needed to examine the effect of native RNA targets of FXR1. However, as the RNA targets of FMR1 and likely FXR1 are quite long (Greenblatt and Spradling, 2018) and impossible to synthesize with current techniques. We speculate that longer RNAs may be required to induce droplet formation *in vitro*. Only isoform E binds G-quadruplex structures (Bechara et al., 2007; Lyons et al., 2017), which promote LLPS (Fay et al., 2017). FXR1 also may have non-specific RNA interactions via the RGG domain. It will therefore be important in the future to understand if specific targets of FXR1 are primarily driving phase separation, and how the different modes of RNA interactions (i.e. KH *versus* RGG) influence droplet assembly.

One of the striking findings in this study is that a longer IDD led to more abundant and functional condensates in myotubes but limits FXR1 LLPS when analyzed in U2OS cells. While in myotubes, isoform E forms more, brighter puncta, in U2OS cells we found that the short isoform A enriches in pellets at lower protein concentrations than the long isoform E and isoform A is more likely to be found in very large and highly spherical droplets. In contrast, in U2OS cells isoform E tends to be in small or irregular droplets, which we suspect are either arrested in fusion or very slow to fuse (**Fig. 5**). Given that S/R-rich sequences can be heavily impacted by PTMs, we hypothesize that the additional sequence provides more residues to enhance modifications that disfavor phase separation in this context.

Interestingly, isoforms A and E aggregate rather than form droplets *in vitro*, which is quite different than results obtained for FMR1 (Tsang et al., 2019). A potential explanation for this difference is that phosphorylation sites of FMR1 are more evenly distributed across the protein rather than strongly concentrated within the IDD as we observed in FXR1. We found that phosphorylation strongly suppressed the FXR1 aggregation phenotype observed *in vitro* with a stronger impact on isoform E whose IDD contains 41 serine residues while isoform A has 26. Phosphorylation has a well-established role in influencing biomolecular condensation (Aumiller and Keating, 2016; Hofweber and Dormann, 2019; Snead and Gladfelter, 2019) and is both rapid and reversible. Phosphorylation has previously been shown to either promote dissolution of granules (Rai et al., 2018; Wippich et al., 2013) or inhibited phase separation (Rai et al., 2018) as we found for FXR1. Interestingly, phosphorylation of the IDD of FMR1, a FXR1 homologue, is required for its interaction with another RNA binding domain Caprin, to tune interactions with RNA. Phosphorylation of FMR1 did not block phase separation suggesting that phosphorylation-dependent inhibition is unique to FXR1 although the authors did not test the full length FMR1 or native FMR1 (Kim et al., 2019). This finding offers a new way of thinking about how splicing regulates PTMs within IDD sequences by altering their length and amino acid composition and their ability to interact with other RBPs and in turn, alter RNA targets.

More surprising was the analysis of proteins missing the C-terminal IDD completely (**Fig. 7E**). A fragment containing only the N-terminus and lacking any predicted IDD robustly formed large, highly spherical droplets. The N-terminal Agenet domains may mediate interactions with other phase-separating proteins to recruit the truncated FXR1. In this case, the Agenet domains would provide multivalency to contribute to the normal LLPS process. The Agenet domains of FMR family proteins bind methylated lysine residues (Adams-Cioaba et al., 2010); therefore, PTMs of binding partners could modulate FXR1 interactions throughout development. Finally, the developmental splicing program may alter which mRNAs are available for FXR1 interaction, and depending on the identity of the bound mRNAs, the IDD role may vary from what we see in U2OS cells.

In summary, we have shown a functional role for FXR1 splicing in development, to promote extension of an IDD relevant for FXR1 condensation with RNA. The unexpected negative effect of a longer IDD on the propensity to phase separate in U2OS cells raises important possible mechanisms for how IDDs may contribute to biomolecular condensates in different cell types. Future work will determine how condensates control translation of associated mRNAs and if FXR1 splice isoforms lead to different targets during tissue identity acquisition and maintenance. In conclusion, we link a developmental splicing program to changes in condensation in a FXR1, a mechanism which may be generalizable to other proteins which undergo phase separation.

## ACKNOWLEDGMENTS

We thank the Gladfelter and Giudice labs, Nancy Kedersha (Brigham and Women’s Hospital), and Silvia Ramos (UNC-Chapel Hill) for critical discussions, Eunice Y. Lee (UNC-Chapel Hill) for technical help, Dr. Stephanie Gupton (UNC-Chapel Hill) for donation of wild type C57Bl/6J mouse embryos, and Marcin Wlizla and NXR (RRID:SCR_013731) for their help in maintaining adult frogs.

## AUTHOR CONTRIBUTIONS

Conceptualization, S.E.R.D, M.E.H., A.S.G., J.G.; Methodology, M.E.H., A.S.G., J.G.; Investigation, J.A.S., E.G.C., R.E.B., C.A.R., S.E.R.D., A.R.V., D.C.J., X.C., S.L., J.C.; Validation, J.A.S., E.G.C., R.E.B., C.A.R., S.E.R.D., X.C., S.M.L., J.G.; Formal Analysis, J.A.S., E.G.C., R.E.B., C.A.R., J.G.; Writing – Original Draft, J.A.S., E.G.C., R.E.B., A.S.G., J.G.; Writing – Review & Editing, J.A.S., E.G.C., R.E.B, C.A.R., S.E.R.D., A.R.V., D.C.J., X.C., S.M.L., J.C., M.E.H., A.S.G., J.G.; Visualization, J.A.S., R.E.B., J.G.; Funding Acquisition, P.A., M.E.H., A.S.G., J.G.; Resources, P.A., A.S.G. and J.G.; Supervision, P.A., M.E.H., A.S.G., J.G.

## METHODS

### Materials

Chemicals and cell culture reagents were obtained from GIBCO, Life Technologies, Thermo Fisher Scientific, and Sigma. C2C12 cells and wild type U2OS cells were purchased from ATCC.

### Mice

FVB/NJ mice were purchased from The Jackson Laboratory. Colonies were derived by mating animals. Mice were handled following the NIH Guidelines for Use and Care of Laboratory Animals that were approved by the Institutional Animal Care and Use Committee (IACUC) at The University of North Carolina at Chapel Hill (UNC-Chapel Hill).

### Mouse tissue isolation

Adult (4-5 months old), and neonatal (P4.5) mice were anesthetized using isoflurane and after cervical dislocation (adult) or decapitation (P4.5) tissues were removed and frozen in liquid nitrogen. Wild type C57Bl/6J mouse embryos at day 15.5 (E15.5) were donated from Dr. Stephanie Gupton’s laboratory (UNC-Chapel Hill). Embryos were rinsed with PBS, the thoracic cavity was opened, and the hearts were dissected. The epidermal layer was then removed and the fore and hind limbs were excised after removing the paws. Embryonic tissue was blotted dry and snap frozen in liquid nitrogen.

### Cell culture

C2C12 undifferentiated cells (myoblasts) were maintained at 37°C under 5% CO_2_ in Dulbecco’s Modified Eagle’s Medium (DMEM) supplemented with 10% fetal bovine serum (FBS) (Gemini Bio-Products), 2 mM glutamine, and 100 units/mL of penicillin, 100 µg/mL streptomycin under low confluency conditions (less than 40%). For differentiation of myoblasts into myotubes, cells were washed with PBS and then cultured in DMEM supplemented with 2% horse serum, 2 mM glutamine, and 100 units/mL of penicillin, 100 µg/mL streptomycin. U2OS cells were maintained at 37°C under 5% CO_2_ in DMEM supplemented with 10% FBS, 100 units/mL of penicillin, and 100 µg/mL streptomycin. For plasmid transfection, cells were transfected at ~60-70% confluency with Lipofectamine 2000 reagent (Invitrogen) following manufacturer’s protocols. In tetracycline (Tet) experiments, 24 h after transfection, cells were induced with Tet for 16 h and subsequently used for the assays.

### Generation of U2OSΔFFF

To create the U2OSΔFFF cell line, *FXR2* was first knocked out using a guide RNA with the sequence 5’-CCC CAT AGG TTC GAG TCG CA-3’ in the U2OS tet-inducible cells. Several clones were selected and FXR2 protein expression was evaluated by immunofluorescence and Western blotting. Clone 6 was selected for future experiments. This cell line was then co-transfected with Cas9 and guide RNAs targeting *FXR1* (5’-TTC CTA GGA ATC TCG TTG GT-3’) and *FMR1* (5’-AAG AGG CGG CAC ATA AGG AT-3’). Clones were selected and screened in a similar manner and finally clone 24 was selected. All loci were sequenced to confirm deletions in the DNA.

### Plasmids

Plasmids used in this study are listed in **Supplementary Table 1**. Restriction enzymes and competent cells were from New England Biolabs (NEB, Ipswich, MA). Oligonucleotides used for plasmid generation were synthesized at Integrated DNA Technologies (Coralville, IA). PCRs were performed using iProof high-fidelity polymerase (Bio-Rad). Sequencing was performed by Genewiz (Morrisville, NC). The sequences of the primers utilized for cloning are detailed in **Supplementary Table 2**.

The mNeonGreen-tagged constructs were created by first making a *pCMV-mNeonGreen* plasmid (pAGB1103). The mNeonGreen ORF was amplified from pAGB1070 with the primers AGO2195 and AGO2196 via PCR. The resulting product was cloned into pAGB879 cut with BglII and SacI (to remove *Fxr1-turboGFP* from the plasmid) using the NEBuilder HiFi DNA assembly reaction (NEB). To make pAGB1111, *Fxr1* isoform E was then amplified with primers AGO2202 and AGO2203 via PCR and cloned via NEBuilder HiFI DNA assembly reaction into pAGB1103 previously cut with NotI and SacII to make an N-terminally tagged construct. To create the C-terminal truncations where the IDD was either truncated back to the RGG (*Fxr1^1-490^*) or completely truncated (*Fxr1^1-374^*), the same approach was taken using different reverse primers to only amplify the first 1122bp of the ORF (pAGB1176, reverse primer was AGO2206) or the first 1470 bp of the ORF (pAGB1177, reverse primer was AGO2207). All of the mutations were created in the tet-inducible mNeonGreen-tagged isoform E construct.

To mutate the KH domains, point mutations were created in the conserved GxxG protein loop required for nucleic acid binding in either the first (*Fxr1^KH1^*), second (*Fxr1^KH2^*), or both domains (*FXR1^KH1/2^*). These mutations should abrogate nucleic acid binding to the KH domain without destabilizing protein structure (Hollingworth et al., 2012; Wang et al., 2015; Wong et al., 2013). These point mutations and deletions were created via site directed mutagenesis using the following primers for each mutation: T236D H237D (KH1) – AGO1869 and AGO1870; K299D N300D (KH2) – AGO1871 and AGO1872. To analyze the effect of loss of the RGG box, this region was completely removed (Δ464-490) to create *Fxr1*^Δ^ using site directed mutagenesis with primers AGO1873 and AGO1874.

To move all mNeonGreen tagged mutations and truncations into the tet-inducible plasmid, pAGB1139 was cut with BamHI and EcoRI so that each insert could be cloned into the plasmid via NEBuilder HiFi DNA assembly reaction (NEB). The primers AGO2350 and AGO2351 were used to amplify the following mNeonGreen-tagged ORFs for assembly into pAGB1139: WT (creating pAGB1162), KH1 (creating pAGB1171), KH2 (creating pAGB1172), KH1/2 (creating pAGB1173), ΔRGG (creating pAGB1174), and ΔRGG KH1/2 (creating pAGB1175). Different reverse primers were used to amplify *mNeonGreen-Fxr1isoE^1-374^* (AGO2374, creating pAGB1166) and *mNeonGreen-Fxr1isoE^1-490^* (AGO2375, creating pAGB1167) for assembly into pAGB1139. To create the mNeonGreen-tagged isoform A construct, isoform A was first created in the pCMV isoform E background (creating pAGB936) using primers AGO1863 and AGO1864 to remove amino acids 569-677 and change amino acids 564-568 from DDSEK to GKRCD, and primers AGO1867 and AGO1868 to remove internal amino acids 308-408. To move this ORF into the tet-inducible system with an N-terminal mNeonGreen tag, mNeonGreen was amplified from pAGB1103 with primers AGO2350 and AGO2486, and *Fxr1isoA* was amplified from pAGB936 with primers AGO2487 and AGO2488. Both inserts were integrated into pAGB1139 cut with BamHI and EcoRI via NEBuilder HiFi DNA assembly reaction (NEB) to create pAGB1189. All constructs and mutations were verified by sequencing.

### Delivery of siRNAs into C2C12 cells

C2C12 cells were seeded into six well plates (8X10^4^ cells/well) in DMEM supplemented with 10% FBS and 2 mM glutamine. The next day, transfections were performed using Lipofectamine RNAiMax transfection reagent (Invitrogen) and Stealth siRNAs (Invitrogen) following manufacturer protocols. The si-RNA sequences were as follows: (i) si-luc (luciferase reporter control, #12935146): GCA CUC UGA UUG ACA AAU ACG AUU (sense) and AAA UCG UAU UUG UCA AUC AGA GUG C (antisense), (ii) si-*Fxr1*-#1 (MSS204455): GCA AUC CAU ACA GCU UAC UUG AUA A (sense) and UUA UCA AGU AAG CUG UAU GGA UUG C (antisense), and (iii) si-*Fxr1*-#2 (MSS204457): GAA GUU GAU GCU UAU GUC CAG AAA U (sense) and AUU UCU GGA CAU AAG CAU CAA CUU C (antisense). The next day, cells were washed with PBS and differentiation was induced for 4-5 days.

### Delivery of MOs into C2C12 cells

C2C12 myoblasts were seeded (8X10^4^ cells/well) in DMEM supplemented with 10% FBS and 2 mM glutamine into six well plates or six well plates with 12 mm coverslips pretreated with 40 µg/mL PureCol (Advanced BioMatrix). The following day, 10-20 µM MOs (Gene Tools, LLC) were delivered using 6-10 µM aqueous Endo-Porter (Gene Tools, LLC) following the suggested protocol from the manufacturer. The MO oligo sequences were as follows: (i) control: CCT CTT ACC TCA GTT ACA ATT TAT A, (ii) *Fxr1* mo-1: AAC AGG CCC ACT CAA GTT ACC TGG C, and (iii) *Fxr1* mo-2: GCA ACT GTG ACT GTT AAA GAT GAG A. The day after delivery, cells were washed with PBS and differentiated in DMEM supplemented with 2% horse serum and 2 mM glutamine for 3-5 days.

### Injection of MOs and sgRNAs in X. tropicalis and X. laevis

*Xenopus* were handled following the NIH Guidelines for Use and Care of Laboratory Animals that were approved by the IACUC at the Marine Biological Laboratory. In **Fig. 1C-D** Xtr.Tg(pax6:GFP;cryga:RFP;act1:RFP)^Papal^ (RRID:NXR_1021) embryos were injected with 20 ng mo-e15i15 or standard control morpholino and analyzed at stage 45. In **Fig. 1E-F** Xla.Tg(actc1:GFP)^Mohun^ (RRID:NXR_007) embryos were injected with 10 ng standard control morpholino, 10 ng mo-i14e15 (targeting both L and S genomes) or co-injected with 1,500 pg Cas9 protein, and 500 pg each sgRNA (T1, T3) and analyzed at stage 35-38. In **Fig. 1G**, one cell at 2-cell stage was injected with 250 pg each sgRNA (T7L, T7S, T8L, T8S) and 1,500 pg Cas9 protein or with 10 ng mo-i14e15. Phenotypes were analyzed at stage 37/38. MO and sgRNA sequence were as follows: (i) *X. tropicalis* mo-e15i15: GTA ACT AAA GTG GGT GGA GCT ATT G, (ii) *X. laevis* mo-i14e15: AAA CTT TGC TGT TGC AGA TGA TAG T, (iii) mo-control: CCT CTT ACC TCA GTT ACA ATT TAT A, (iv) T1 sgRNA: CCC CTG AAG CGA CGC CTG CGG, (v) T3 sgRNA: GGC AGA AGA TAG ACA GCC AGG, (vi) T7L sgRNA: ACT GCA GGT TGC AAA CAT ATT GG, (vii) T7S sgRNA: TAC TAT TGC TAG TCT TCA AGA GG, (viii) T8L sgRNA: TCA GGA CAA TGG TTC TTA AAT GG, and (ix) T8S sgRNA: AAA TGA TAG AAT GTC TAG TGT GG. All sgRNA include a PAM sequence.

### RNA extraction

Frozen mouse tissues were homogenized using 1.4 mm ceramic beads (Lysing Matrix D) and a Precellys-24 homogenizer (Bertin Instruments) with the following setting: 6,500 r.p.m. for 20 s intervals until complete homogenization. RNA was then extracted following TriZol (Invitrogen) manufacturer’s protocols. For cell culture samples, TriZol reagent was added to the plates followed by RNA extraction per the manufacturer suggested protocol. For *Xenopus* experiments, RNA was extracted from whole embryos (7 embryos per sample) using TriZol following manufacturer protocol. RNA concentration was measured with a NanoDrop Lite Spectrophotometer (ND-LITE, ThermoScientific).

### cDNA synthesis

Commercial human RNA samples (**Supplementary Table 3**) or RNA extracted from mouse tissues, *Xenopus* embryos, or cells was utilized to prepare cDNA using the High-capacity cDNA reverse transcription kit (Applied Biosystems) or the SuperScript IV kit (Invitrogen). The program was as follows: (i) 25°C for 10 min, (ii) 37°C for 120 min, (iii) 85°C for 5 min, (iv) 4°C pause.

### Analysis of Fxr1 alternative splicing

The synthesized cDNA was used to perform PCR assays with the mouse (0.5 µM), *X. tropicalis* (0.4 µM), or human primers (0.5 µM) targeting the constitutive exons flanking the alternative regions (**Supplementary Table 4**). PCRs for C2C12 cells, mouse tissues, and human samples were performed using GoTaq green master mix (Promega) and the following amplification conditions: (i) 95°C for 1 min 15 sec, (ii) 28 cycles of 95°C for 45 s, 57°C for 45 s, 72°C for 1 min, (iii) 72°C for 10 min, (iv) 4°C pause. PCR products were separated by 6% polyacrylamide gel electrophoresis in TBE buffer (89 mM Tris, 89 mM boric acid, 2.5 mM EDTA, pH 8.3) for 3-4 h at 165 V. After electrophoresis, the gels were stained with 0.4 µg/mL ethidium bromide in water for 10 min and visualized using the ChemiDoc^TM^ XRS+ (Biorad) imaging system. Quantification of gels was performed by densitometry using Image Lab^TM^ 6.0.1 software for analysis (Biorad). In *X. tropicalis* experiments, PCRs were performed using Taq Polymerase (NEB standard protocol) and the following amplification conditions: (i) 95°C for 1 min, (ii) 27 cycles of 90°C for 30 s, 60°C for 30 s, 72°C for 16 s, (iii) 72°C for 5 min, (iv) 4°C pause. PCR products were separated by 2% agarose gel electrophoresis for 40 min at 135 V.

### Real-time PCR (qPCR)

A 20 µL reaction with TaqMan Fast Advanced Master Mix (Applied Biosystems) was used to quantify 50-100 ng cDNA with TaqMan probes for *Fxr1* (Applied Biosystems, Mm00484523-m1, amplicon size 73 bp), and *Hmbs* (Applied Biosystems, Mm01143545-m1, amplicon size 81 bp) transcripts. An Applied Biosystems StepOnePlus Real-Time PCR System was used and the thermal-cycling profile was as follows: (i) 50°C for 2 min, (ii) 95°C for 20 s, (iii) 40 cycles of 95°C for 1 s, 60°C for 20 s. The CT (cycle threshold) values of quantified transcripts were normalized to that of a reference gene (*Hmbs*) from the same sample.

### Protein extraction from mouse tissues

Frozen tissues were homogenized with 1.4 mm ceramic beads (Lysing Matrix D, MP Biomedicals) in a Hepes/sucrose buffer (10 mM Hepes, 320 mM sucrose, 1% sodium dodecyl sulfate) supplemented with protease and phosphatase inhibitors (Sigma Aldrich). The Precellys-24 homogenizer was utilized at 6,500 r.p.m. for 20 s intervals until complete tissue homogenization. Samples were then sonicated in an ice water bath for 3 min (30 s bursts) at 75 V. Lysates were collected after spinning at 14,000 rpm for 10 min at 4°C, and the protein concentration was measured using the Pierce BCA protein assay kit (Thermo Fisher Scientific).

### Protein extraction from cell culture

Cells were lysed in RIPA buffer (1% Triton X-100, 0.1% SDS, 0.5% sodium deoxycholate, 150 mM NaCl, 50 mM Tris, 5 mM EDTA, pH 8.0) supplemented with protease and phosphatase inhibitors (Roche). Cell lysates were sonicated for 3 min (30 s bursts) at 75 V. After spinning at 14,000 r.p.m. for 10 min at 4°C., supernatants were collected and protein concentration was measured using the Pierce BCA protein assay kit. To analyze insoluble aggregates by Western blot, C2C12 or U2OS cells were lysed in NETN lysis buffer (250 mM NaCl, 5 mM EDTA pH 8.0, 50 mM Tris-HCl pH 8.0, 0.5% NP-40) supplemented with protease inhibitors for 30 min on ice. Lysates were clarified by centrifugation for 10 min at 13,000 r.p.m. at 4°C. The soluble lysate was separated from the pellet and the pellet was resuspended in 50 µL of lysis buffer. Equivalent fractions were run on an SDS-PAGE gel for analysis via Western blotting.

### Western blot assays

For **Figs. 3C, 4E, S2C, S3A**, protein samples (25 µg) prepared in loading buffer (62.5 mM Tris-HCl, 10% glycerol, 2% SDS, 0.02% bromophenol blue, 143 mM beta-mercaptoethanol, 5 mM dithiothreitol, pH 6.8) were boiled for 5 min and then loaded into 10% Mini-Protean TGX (Tris-Glycine eXtended) Stain-Free precast Gels (Biorad). Electrophoresis was performed in a buffer containing 25 mM Tris, 190 mM glycine, and 0.1% w/v SDS (pH 8.3) for 2 h at 120 V. After electrophoresis, the gels were exposed to UV light for 2.5 min to activate TGX within the gel or counter stained with Ponceau stain solution for total protein visualization. Proteins were transferred onto 0.45 µm PVDF Immobilon-FL membranes (Millipore Sigma) and visualized using the ChemiDoc XRS+ (Biorad) imaging system. Membranes were blocked with 5% non-fat dried milk in TBST buffer (19 mM Tris, 2.7 mM KCl, 137 mM NaCl, 0.1% Tween20, pH 7.4) for 1 h, washed, and incubated overnight at 4°C with primary antibodies diluted in 5% bovine serum albumin (BSA) in TBST. Primary antibodies were as follows: (i) rabbit monoclonal [EPR7932] anti-FXR1 (Abcam, ab129089) diluted 1:1,000, (ii) mouse monoclonal anti-GAPDH (Santa Cruz Biotechnology, sc365062) diluted 1:500, (iii) mouse monoclonal anti-alpha tubulin (Santa Cruz Biotechnology, sc32293) diluted 1:500, and (iv) rabbit polyclonal anti-beta tubulin (Abcam, ab6046) diluted 1:1,000. The next day, the membranes were washed three times (10 min each) with TBST and then incubated for 1-1.5 h at room temperature in the darkness with a goat polyclonal anti-rabbit IgG (H+L) DyLight 800 4X PEG (Invitrogen, SA5-35571) diluted (1:10,000) in 5% BSA in TBST. Membranes were washed three times (10 min each) with TBST and fluorescent signal was detected using the Odyssey CLx Blot Imager (Li-Cor). For **Fig. 4**, **5** and **7**, Western blots were performed as above with the following differences. SDS-PAGE gels were transferred to a nitrocellulose membrane and subsequently blocked for 2 days in 5% milk in 50 mM Tris, 150 mM NaCl, 0.1% Tween20. Membranes were washed and incubated overnight at 4°C with primary antibodies. Primary antibodies were as follows: (i) rabbit monoclonal [EPR7932] anti-FXR1 (Abcam, ab129089) diluted 1:1,000, and (ii) rabbit anti-CDC2 (Santa Cruz Biotechnology) diluted 1:1,000. The membranes were then washed three times (10 min, 5 min, 5 min) and incubated for 1 h at room temperature with goat anti-rabbit IgG (H+L) (BioRad). Membranes were washed and signal was detected using the ChemiDoc XRS+ (Biorad) imaging system.

### Immunofluorescence assays

Cells in plates or coverslips were washed three times with PBS and fixed in 4% paraformaldehyde (Electron Microscopy Sciences) in PBS for 20 min at room temperature. Cells were washed three times in PBS, incubated in blocking solution (137 mM NaCl, 2.7 mM KCl, 9.6 mM Na_2_HPO_4_, 1.5 mM KH_2_PO_4_, 1% BSA, 0.3% Triton X-100, pH 7.4) for 1 h at room temperature, and then with the primary antibodies diluted in blocking solution overnight at 4°C. Primary antibodies were as follows: (i) rabbit monoclonal [EPR7932] anti-FXR1 (Abcam, ab129089) diluted 1:250, and (ii) mouse monoclonal [B-5] anti-MYH (Santa Cruz Biotechnology, sc-376157) diluted 1:50. The next day, cells were washed three times (10 min each) with PBS and then incubated with the appropriate secondary antibody diluted in blocking solution for 1 h at room temperature in the darkness. Secondary antibodies were as follows: (i) rabbit polyclonal anti-mouse IgG (H+L) Alexa Fluor 488 (Invitrogen, A21204) diluted 1:500, and (ii) goat polyclonal anti-rabbit IgG (H+L) Alexa Fluor 488 (Invitrogen, A11008) diluted 1:250. Cells were washed three times (10 min each) with PBS and then stained with 2 µM DAPI in PBS for 5 min at room temperature in the darkness. Cells were washed three times (5 min each) in PBS and imaged by confocal microscopy in PBS. When cells were stained with phalloidin, incubation with Alexa Fluor 647 Phalloidin (Invitrogen, A22287, dilution 1:40) was performed concurrently with the secondary antibody.

### Confocal microscopy for cells

For imaging in **Fig. S3**, confocal microscopy was performed in the Hooker Imaging Core (UNC-Chapel Hill) using a Zeiss LSM 880 confocal microscope with a 10X Plan Apochromat objective (0.45 WD). Excitation parameters were as follows: argon multiline laser at 488 nm (alexa fluor 488) (2% power), a 405 nm diode at 30 mW (DAPI) (2% power), or a Helium-Neon laser at 633 nm (2% power). Emission filters were set up as follows: 490-615 nm (alexa fluor 488), 410-514 nm (DAPI), and 638-747 nm (alexa fluor 647). For imaging in **Fig. 4C**, live cells were imaged on a Quorum WaveFX spinning disk confocal system equipped with a Plan Apo 60X 1.4 NA objective. Images were acquired on an electron multiplying (EM)-CCD device (C9100-13, Hamamatsu) driven by Metamorph software version 7.8.12.0 (Molecular Devices, Sunnyvale, CA). A 488 nm laser was used for GFP excitation using 20% laser power, 500 ms exposure, 200 gain. Cells were imaged every 15 s through the complete depth of the cell with 0.5 µm inter-plane spacing. For **Figs. 4B**, **4G**, **5**, and **7**, fixed or live cells were imaged on Yokogawa CSU-W1 spinning disk confocal system equipped with a VC Plan Apo 60X 1.49 NA oil immersion objective. Images were acquired using a sCMOS 85% QE camera (Photometrics) driven by Nikon Elements Software. For **Fig. 4B** and **4G** the following parameters were used: 488 nm laser (alexa fluor 488) at 25% power and 200 ms exposure, 405 nm laser (Hoechst) at 15% power and 100 ms exposure, 640 nm laser (alexa fluor 647 Phalloidin) at 50% power and 100 ms exposure. For **Figs. 5** and **7** the following parameters were used: 488 nm laser (mNeonGreen) at 25% power and 100 ms exposure, 405 nm laser (Hoechst) at 15% power and 100 ms exposure.

### Microscopy in Xenopus experiments

Images of *Xenopus* tadpoles in **Fig. 1** were acquired on a Zeiss SteREO Discovery.V12 microscope with an Achromat S 1.25X objective and a mounted Zeiss AxioCam MRc camera controlled with the ZEN 2 lite (blue edition) software version 2.0.0.0. The filter blocks 45 Texas Red/FRFP and 57 GFP LP were used to acquire red or green fluorescent images respectively. Prior to imaging, the tadpoles were anesthetized until complete loss of mobility using 0.1 % Tricaine-S (Western Chemical, Inc.) buffered to pH 7.4 with NaHCO_3_.

### Protein purification

FXR1 isoforms A and E were cloned into the HIS-GB1 vector and Sanger sequenced to confirm insert. Plasmids were transformed into BL21 bacteria. 2XYT media containing kanamycin was inoculated with a colony of freshly transformed Bl21 (DE3). Cultures were grown at 37°C overnight with shaking. The next morning, the starter culture was diluted 1:60 and was incubated at 37°C for 2 h, and then at 30°C until culture reached an optical density (OD) 0.6-0.7. The culture was induced with 1 mM IPTG and then incubated at 30°C for 6 h. Cells were pelleted at 10,000 × r.p.m. for 5 min and pellets were resuspended in 1.5 M KCl, 50 mM HEPES pH 7.4, 20 mM imidazole, 5 mM beta-mercaptoethanol, 1.5 mL 10 mg/mL lysozyme, and protease inhibitor for 30 min at 4°C. Samples were then sonicated (10 s on, 2 min off, 10 s on) until homogenous. Samples were centrifuged at 20,000 r.p.m. for 30 min and the supernatant was passed through a 0.44 µm filter using a 60 mL syringe. FXR1 was purified from the supernatant using a Ni/Co-NTA column following standard protocols for Whi3 purification (Langdon et al., 2018). Purified proteins were analyzed by SDS-PAGE and Coomassie staining.

### Casein kinase II *in vitro* phosphorylation assay

*In vitro* phosphorylation reactions were carried out according to the manufactures instructions (NEB, P6010S) using 0.9 nanomoles of protein with control phosphorylation performed in the absence of ATP with all other reaction components. Reactions were incubated for 50 min at 30°C. In the last 10 min of incubation 60 µM Atto 488 dye (Sigma, 41051-1MG-F) was added to the reaction. Following incubation and labeling (1 h total), samples were buffer exchanged using CENTRI SPIN-20 columns (Princeton Separations, CS-201) according to the manufacturers instructions to remove the buffer, excess dye, and ATP. Protein concentration was requantified using a nanodrop. Phosphorylation was confirmed by gel electrophoresis followed by Coomassie staining.

### In vitro transcription

Cy5 labeled luciferase RNA was produced according to the manufacturers instructions (NEB, E2040S) labeled with 6% Cy5-UTP (PA53026, Milipore sigma). Reaction was incubated at 37°C overnight and precipitated using 2.5 M LiCl. Pellets were washed three times with 70% ethanol in DEPC treated water. RNA samples were eluted in DEPC treated water and quantified using a nanodrop. *In vitro* transcribed RNA (~5 µg) was analyzed by agarose gel electrophoresis to confirm the presence of a single band with the appropriate size.

### In vitro assays

10 nM of luciferase RNA and 2 µM or 4 µM FXR1 isoform A and E (Atto 488 labeled) were mixed in droplet buffer (150 mM KCL, 20 mM HEPES, 1 mM BME, pH 7.4) and added to 30 mg/mL fatty acid free BSA (Sigma, A8806) in droplet buffer blocked chamber slides (Grace Bio-Labs, 2024-03) and incubated at room temperature for 3 h prior to imaging.

### Image processing and quantitative analysis

Processing of microscopy images and quantitative analysis were performed using Fiji software (Schindelin et al., 2012; Schneider et al., 2012) or ImageJ software (imagej.nih.gov). Images were identically contrasted with respect to black and white values within each figure. The fusion index was estimated as the ratio between the number of nuclei within myosin heavy chain (MYH) positive cells with >2 nuclei and the total number of nuclei. To quantify the coefficient of variation (CV) in **Fig. 4**, a region of interest was first drawn around the most focused z-stack for each cell with >2 nuclei excluding the nuclei. The mean gray value was acquired in the FXR1 channel by calculating the sum of the gray values of all the pixels in the region of interest divided by the number of pixels. This value was then background subtracted. The standard deviation of the gray values for each cell was then quantified. The CV was then calculated by dividing the standard deviation of a cell by its background subtracted mean gray value. For droplet formation analysis, cells were visually scored for either having droplets or not. For morphology quantification, cells were visually scored for the three morphology categories observed. For in vitro assays, images were quantified using Fiji software as follows: Protein signal (488) was used to threshold to white level of 137 (a.u.) and the Analyze Particle plugin was utilized to define the area of the particles in the image. These regions of interest (ROIs) were then used to extract the RNA and protein fluorescent signal. Three images from three replicates were used for quantification.

### Software

Protein disorder was predicted using the IUPRED algorithm (Dosztányi et al., 2005a, 2005b). The amino acid composition of the C-terminal IDD of FXR1 was determined using IDD Navigator with IUPRED disordered domain prediction (Patil et al., 2012) along with the ProtParam tool (https://web.expasy.org/protparam/) (Gasteiger et al., 2005). For phosphorylation prediction, the sequences of the HIS-GB1-FXR1 A/E, or FMR1 (uniprot) were analyzed using the NETphos 3.1 predictor (serine phosphorylation) and IUPRED (disorder).

### Statistics

All data are presented as the mean ± s.e.m. The p-values were estimated by a Student’s t-test (two tailed) or one-way ANOVA with Bonferroni test for multiple comparisons. p≤0.05 was considered statistically significant.

