## Supplemental Material for "Regulation of FXR1 by alternative splicing is required for muscle development and controls liquid-like condensates in muscle cells"

### SUPPLEMENTAL FIGURES

*Xenopus laevis*
T1 T3 PAM Mismatch

FXR1 5S Sequence
WT
GTAGCCGCAGGCGTCGCTTCAGGGGTCAGGCAGAAGATAGACAGCCAGg
N1
GTAGC------------TTCAGGGGTCACGCCGAGGATCA--AGNNCCACG
N2
GTAGCCGC----GTCGCTNCTGGC-GGGG--A-GCAGAA-A-------CNNGg
N3
GTAGCCGCAGGCG--G-----G-----ANGCNG-AGNA-A-A-------
N4
GTAGCCGCAG---TCGCTTCAGGGGTCAGGCA-AA---AGATAGCCAGg
N5
GTAGCCGC----GNCNCTTCTGGGGTCAGGCAGAAGATAGACAGACAGg

FXR1 5L Sequence
WT
GTAGCCGCAGGCGTCGCTTCAGGGGTCAGGCAGAAGATAGACAGCCAGg
N1
GTAGCCGCAGG------------GGTCAGGCAGAAGAT--ANAG--AGg
N2
GTAGCCGC-------G-----GGGGTCATGCAGAGGATAG---G-CAGg
N3
GTAGCCGCAGG-G--G--TCAGG---C-GGAAG-AGA-A-ACGGCTA-G
N4
GTAGCCGCANGCGTCTCGCTTCAGGGGTCANG--A-AAAATA-------
N5
GTAGCCGCA-NC--CG-TTCCTGGGNG---GG--G--------CAG---

### SUPPLEMENTAL FIGURE S1

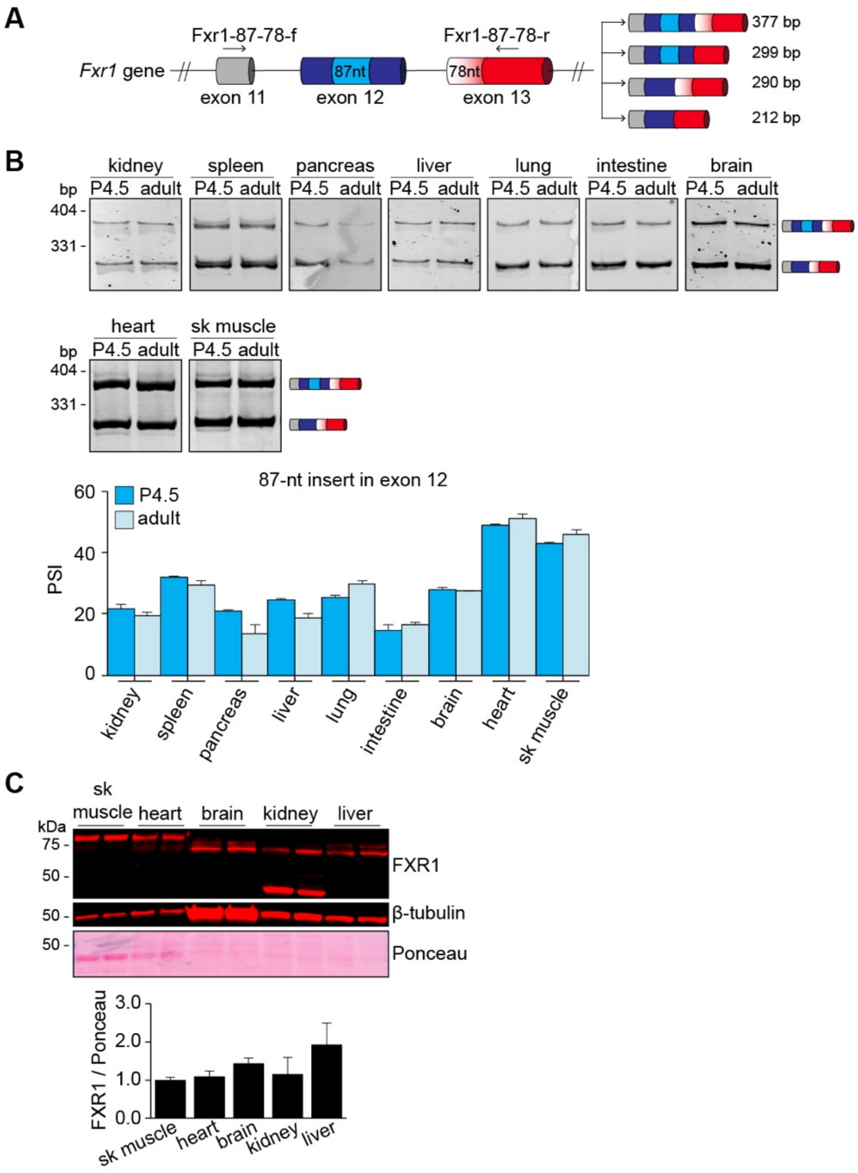


### SUPPLEMENTAL FIGURE S2

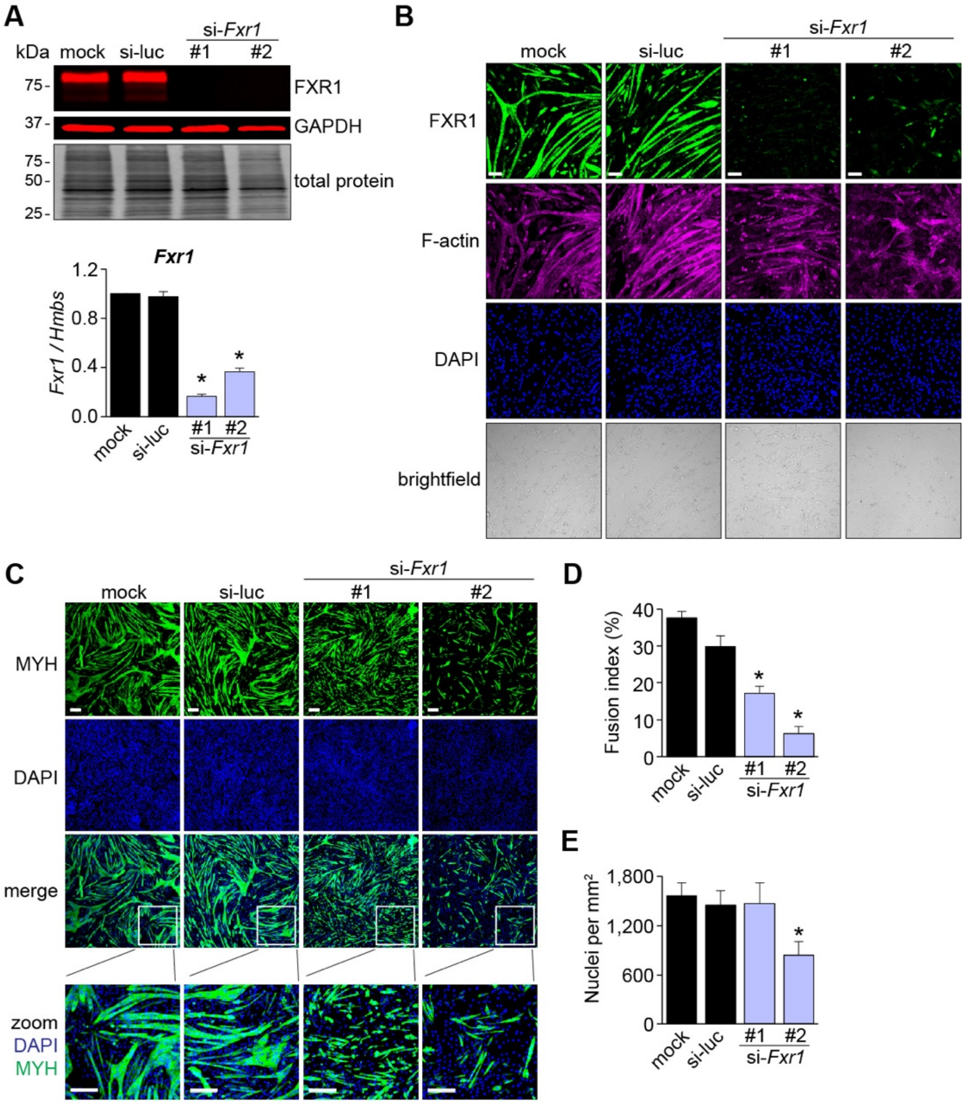


**SUPPLEMENTAL FIGURE S3**

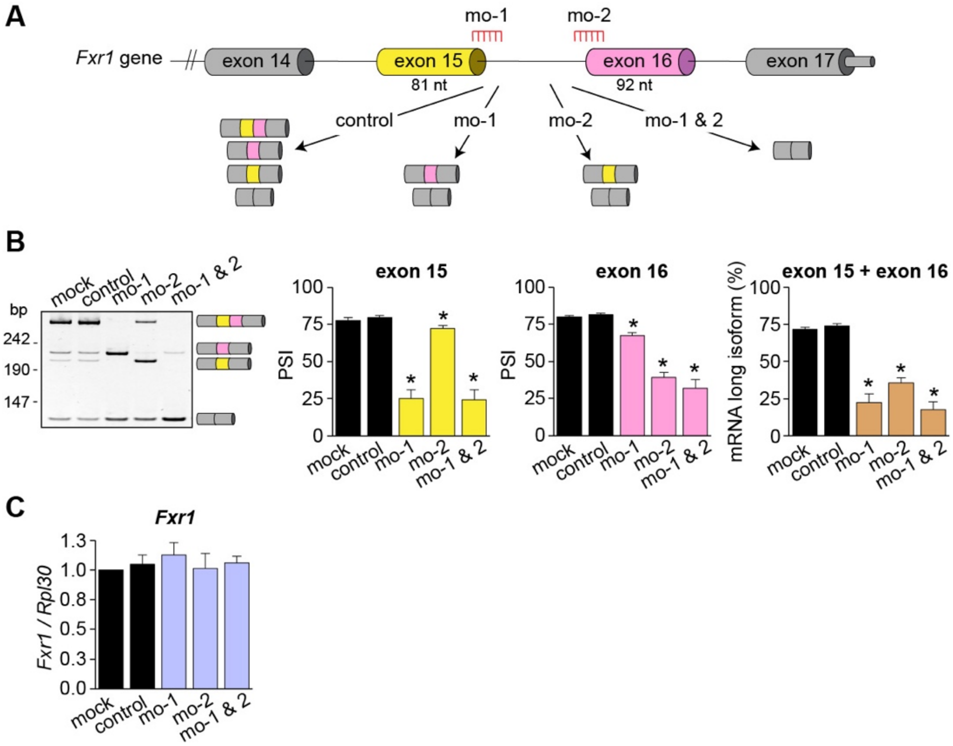


**SUPPLEMENTAL FIGURE S4**

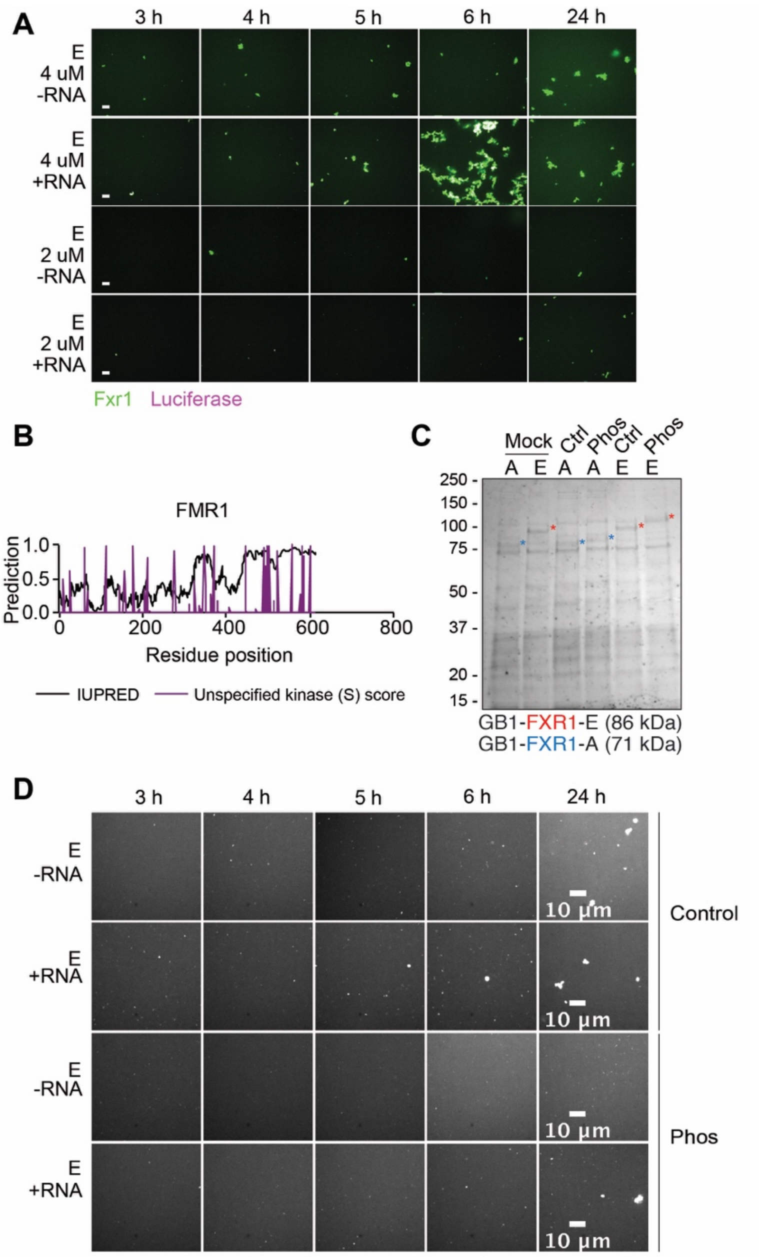


**SUPPLEMENTAL FIGURE S5**

#### SUPPLEMENTAL FIGURE LEGENDS

**Supplemental Figure S1. Sequences of *fxr1* exon 15 from sgRNA treated embryos**. Sequences of *fxr1.S fxr1.L* from five randomly selected *X. laevis* embryos injected with the sgRNA T1 (green) and T3 (underlined). sgRNA injected embryos had deletions (dashed) uncallable nucleotides (N) and mismatches (red) compared to reference sequences. Deletions roughly correspond to 3-4 nucleotides upstream of the PAM sequences (magenta). **Supplemental Figure S1** is linked to **Figure 1E-F**.

**Supplemental Figure S2. Alternative splicing events in exons 12 and 13 of *Fxr1* pre-mRNA and expression of FXR1 protein isoforms during tissue development. A.** Scheme of part of *Fxr1* gene showing the alternative splicing events in exons 12 (insert of 87 nt) and exon 13 (alternative splice site leading to the inclusion or skipping of a 78-nt region). To evaluate the splicing patterns by RT-PCR, the primers designed to target the constitutive flanking exons were Fxr1-87-78-f (forward) and Fxr1-87-78-r (reverse). **B.** Different tissues were collected from neonatal (postnatal day 4.5, P4.5) and adult (4 months) mice. Alternative splicing was evaluated by RT-PCR and quantified by densitometry. *N*=3-4 (P4.5 tissues) samples from 2-9 pooled neonates each depending on tissue type, *N*=4 (adult tissues) animals. **C.** Adult mouse tissues were evaluated by Western blots utilizing a monoclonal antibody against the N-terminus of FXR1 protein that thus recognizes all of the splice variants. *N*=2 animals. Results: mean ± s.e.m. *$p \leq 0.05$ (one-way ANOVA test with Bonferroni correction for multiple comparisons). bp: base pairs, sk. muscle: skeletal muscle. **Supplemental Figure S2** is linked to **Figure 2**.

**Supplemental Figure S3. FXR1 is required for myogenesis.** C2C12 myoblasts were transfected with two different si-RNAs targeting *Fxr1* (#1, #2) or a control siRNA (si-luc) and the next day differentiation was induced. **A.** Protein and RNA were collected at differentiation day 4 and FXR1 depletion was confirmed by Western blot and qPCR assays. **B.** Cells were evaluated by immunofluorescence to confirm knock down. Scale bars 100 µm. **C-D.** Myosin heavy chain (MYH) expression was evaluated by immunofluorescence (**C**), and the fusion index was estimated (**D**). **E.** The number of nuclei per field of view was estimated from the microscopy images. Results: mean ±

s.e.m. *p≤0.05 *versus* mock and si-luc (Student T-test), *N*=4-8 independent experiments. Scale bars 200 μm. **Supplemental Figure S3** is linked to **Figure 3**.

**Supplemental Figure S4. Modulation of endogenous *Fxr1* splicing in C2C12 cells using morpholinos.** Morpholinos (MOs) were delivered in undifferentiated myoblasts and the next day cells were differentiated for four days. **A.** Scheme of the MOs used to target exons 15 (mo-1) and 16 (mo-2). Control denotes cells treated with a MO control. **B.** Efficiency of MO action was evaluated by RT-PCR. **C.** Total *Fxr1* mRNA levels were evaluated by qPCR. Results: mean ± s.e.m. *p≤0.05 versus mock and control (Student T-test), *N*=3-8 independent experiments. **Supplemental Figure S4** is linked to **Figure 4**.

**Supplemental Figure S5. RNA addition and protein phosphorylation influence FXR1 aggregation *in vitro*. A.** Luciferase RNA accelerates isoform E aggregation. Representive images for either 4 μM or 2 μM isoform E with or without luciferase RNA after 3, 4, 5, 6, and 24 h. Scale bars 10 μm. **B.** FMR1 Predicted disorder (IUPRED) and unspecified kinase score for serine are marked for each amino acid. Predicted phosphorylated serines are uniformly distributed across the protein. **C.** Coomassie staining of the *in vitro* purified FXR1 isoforms A and E. Phosphorylation results in a band shift with respect to the control or mock treated samples. **D.** Representive images of 2 μM of FXR1 isoform E (shown in white) with or without RNA and with or without phosphorylation (phos) at 3, 4, 5, 6, and 24 h. Scale bars 10 μm. **Supplemental Figure S5** is linked to **Figure 6**.

**SUPPLEMENTARY TABLES**

| Strain | Genotype | Parent | Reference |
|---|---|---|---|
| AGB828 | $P_{CMV}$-*eGFP-FXR1* (HeLa isoform A) *KAN/NeoR* $Amp^R$ | pEGFP-C1 | Generous gift from Sumara group, IGBMC |
| AGB879 | $P_{CMV}$-*Fxr1isoE-tGFP* $Amp^R$ | pCMV6-AC-GFP | Origene |
| AGB1070 | *mNeonGreen* $Amp^R$ | pNCS | Allele Biotechnology |
| AGB1103 | $P_{CMV}$-*mNeonGreen-Fxr1isoE* $Amp^R$ | pCMV6-AC-GFP | This study |
| AGB1111 | $P_{CMV}$-*mNeonGreen* $Amp^R$ | pAGB1103 | This study |
| AGB1139 | $P_{CMV}$*2xTetO*$_2$ $Amp^R$ $Zeo^R$ | pcDNA4/TO | This study |
| AGB1162 | $P_{CMV}$*2xTetO*$_2$-*mNeonGreen-Fxr1isoE* $Amp^R$ $Zeo^R$ | pcDNA4/TO | This study |
| AGB1189 | $P_{CMV}$*2xTetO*$_2$-*mNeonGreen-Fxr1isoA* $Amp^R$ $Zeo^R$ | pcDNA4/TO | This study |
| AGB1171 | $P_{CMV}$*2xTetO*$_2$-*mNeonGreen-Fxr1isoE*$^{T236D\ K237D}$ $Amp^R$ $Zeo^R$ | pcDNA4/TO | This study |
| AGB1172 | $P_{CMV}$*2xTetO*$_2$-*mNeonGreen-Fxr1isoE*$^{K299D\ N300D}$ $Amp^R$ $Zeo^R$ | pcDNA4/TO | This study |
| AGB1173 | $P_{CMV}$*2xTetO*$_2$-*mNeonGreen-Fxr1isoE*$^{T236D\ K237D\ K299D\ N300D}$ $Amp^R$ $Zeo^R$ | pcDNA4/TO | This study |
| AGB1174 | $P_{CMV}$*2xTetO*$_2$-*mNeonGreen-Fxr1isoE*$^{\Delta 464\text{-}490}$ $Amp^R$ $Zeo^R$ | pcDNA4/TO | This study |
| AGB1175 | $P_{CMV}$*2xTetO*$_2$-*mNeonGreen-Fxr1isoE* $^{\Delta 464\text{-}490\ T236D\ K237D\ K299D\ N300D}$ $Amp^R$ $Zeo^R$ | pcDNA4/TO | This study |
| AGB1176 | $P_{CMV}$*2xTetO*$_2$-*mNeonGreen-Fxr1isoE*$^{1\text{-}374}$ $Amp^R$ $Zeo^R$ | pcDNA4/TO | This study |
| AGB1177 | $P_{CMV}$*2xTetO*$_2$-*mNeonGreen-Fxr1isoE*$^{1\text{-}490}$ $Amp^R$ $Zeo^R$ | pcDNA4/TO | This study |

**Supplementary Table 1.** Plasmids used in this study

| Primer name | Primer sequence (5' to 3') |
|---|---|
| AGO2195 | actggatccggtaccgaggaggagcaggtgcaggtgcaggtgCTCGAGATGGTGAGCAAG |
| AGO2196 | caggaaacagctatgaccgcggagcaggtgcaggtgcaggtgGCGGCCGCTTACTTGTAC |
| AGO2202 | gcatggacgagctgtacaaggcgcgtacgcggccgctcgagATGGCGGAGCTGACGGTG |
| AGO2203 | caggaaacagctatgaccgctcaTGAAACACCATTCAGGACTGCTG |
| AGO2206 | caggaaacagctatgaccgctcaCTGCTCATCAATCTGCAGACGTTC |
| AGO2207 | caggaaacagctatgaccgctcaGCCACCACGTGGTCCACC |
| AGO1869 | GGGCCTGGCGATAGGAGACGATGGCAGTAACATACAGCAAGC |
| AGO1870 | GCTTGCTGTATGTTACTGCCATCGTCTCCTATCGCCAGGCCC |
| AGO1871 | CCCAGGAATCTTGTTGGAAAAGTAATTGGAGACGATGGCAAAGTTATTCAAGAAATAGTGG |
| AGO1872 | CCACTATTTCTTGAATAACTTTGCCATCGTCTCCAATTACTTTTCCAACAAGATTCCTGGG |
| AGO1873 | GATGATCGAGAGACTCGACATAAATCCTCCATCAGTTCTGTGC |
| AGO1874 | GCACAGAACTGATGGAGGATTTATGTCGAGTCTCTCGATCATC |
| AGO2350 | cgtttaaacttaagcttggtaccgagctcgATGGTGAGCAAGGGCGAG |
| AGO2351 | agcggccgccactgtgctggatatctgcagTCATGAAACACCATTCAGGACTG |
| AGO2374 | agcggccgccactgtgctggatatctgcagTCACTGCTCATCAATCTGCAG |
| AGO2375 | agcggccgccactgtgctggatatctgcagTCAGCCACCACGTGGTCC |
| AGO1863 | CCTCAGTTAATGAGAATGGGCTAGGCAAGAGGTGCGACACGCGTACGCGGCCGCTCGAGATGGAGAGC |
| AGO1864 | GCTCTCCATCTCGAGCGGCCGCGTACGCGTGTCGCACCTCTTGCCTAGCCCATTCTCATTAACTGAGG |
| AGO1867 | GCAGCTGCGACAGATTGGTTCTAGGTCTTATAGTGGAAGAGG |
| AGO1868 | CCTCTTCCACTATAAGACCTAGAACCAATCTGTCGCAGCTGC |
| AGO2486 | gcggccgcgtacgcgcCTTGTACAGCTCGTCCATGC |
| AGO2487 | gacgagctgtacaaggcgcgtacgcggccgctcgagATGGCGGAGCTGACGGTG |
| AGO2488 | agcggccgccactgtgctggatatctgcagCTAGTCGCACCTCTTGCCTAG |

**Supplementary Table 2.** Sequence of the primers (IDT) used for plasmid generation. Upper case denotes sequences that will directly base pair with the plasmid being amplified while lower case bases are overhangs for NEBuilder HiFi DNA assembly reaction (NEB).

| Tissue | Sex | Age of individual | Company and catalog number | Lot number |
|---|---|---|---|---|
| heart | male | 21 years | Amsbio (R1234139-50) | B707146 |
| heart | male | 51 years | Cell Applications Inc (1H30-50) | 1369 |
| heart | male | 30-39 years | TakaRa (636532) | 1610293A |
| skeletal muscle | female | 48 years | Amsbio (R1234171-50) | B207200 |
| skeletal muscle | male | 58 years | Cell Applications Inc (1H60-50) | 1376 |
| skeletal muscle | male, female | 20-68 years | TakaRa (636534) | 1609179A |
| heart | female | 22-week of gestation | Cell Applications Inc (1F30-50) | 2739 |
| heart | male | 20-21 weeks of gestation | Agilent (540165) | 0006262824 |
| skeletal muscle | female | 22-week of gestation | Cell Applications Inc (1F60-50) | 2739 |
| skeletal muscle | female | 18 weeks of gestation | Agilent (540181) | 0006260887 |

**Supplementary Table 3.** Commercially available human RNA samples.

| Species | Primer name | Primer sequence (5' to 3') | Analyzed region | Expected bands (bp) |
|---|---|---|---|---|
| mouse | Fxr1-81-92-f | TGCTGTTCTGATGGATGGAC | exon 15 (81 nt) and exon 16 (92 nt) | 300<br>219<br>208<br>217 |
| mouse | Fxr1-81-92-r | GAAGCGCTAGTTGGACCATT | | |

| | | | | |
|---|---|---|---|---|
| mouse | Fxr1-87-78-f | AGAAAGCATTGGGAATGTGC | 87-nt insert in exon 12 and alternative 3' splice site in exon 13 | 377<br>299<br>290<br>212 |
| mouse | Fxr1-87-78-r | TGTCTCGCTGATGTCGAGTC | | |
| human | FXR1-81-92-f | TGCTGTTCTGATGGATGGAA | exon 15 (81 nt) and exon 16 (92 nt) | 298<br>217<br>206<br>125 |
| human | FXR1-81-92-r | AGCACTAGTTGGGCCGTTTA | | |
| *X.tropicalis* | fxr1-F1 | TCTCACCACAACACAAACCG | exon 15 (81 nt) | 266<br>185 |
| *X.tropicalis* | fxr1-R1 | GCGGAATCTACCTAAAGAACC | | |
| *X.tropicalis* | eef1a1-F | CCCCTCTTGGTCGTTTTGCTGTCC | Forward primer spans exons 7-8 junction; reverse is end of coding sequence in exon 8. | 135 |
| *X.tropicalis* | eef1a1-R | TTGCCTTTCTGTGCTTTCTGAGCAG | | |

**Supplementary Table 4.** Sequence of the primers (Sigma, IDT) utilized to evaluate *FXR1* alternative splicing in mouse, human, and frog samples. eef1a1 was used as a loading control. bp: base pairs, nt: nucleotides.
